# Combinatorial quantification of 5mC and 5hmC at individual CpG dyads and the transcriptome in single cells reveals modulators of DNA methylation maintenance fidelity

**DOI:** 10.1101/2023.05.06.539708

**Authors:** Alex Chialastri, Saumya Sarkar, Elizabeth E. Schauer, Shyl Lamba, Siddharth S. Dey

**Author notes:** Correspondence to: Siddharth S. Dey.

## Abstract

Transmission of 5-methylcytosine (5mC) from one cell generation to the next plays a key role in regulating cellular identity in mammalian development and diseases. While recent work has shown that the activity of DNMT1, the protein responsible for the stable inheritance of 5mC from mother to daughter cells, is imprecise; it remains unclear how the fidelity of DNMT1 is tuned in different genomic and cell state contexts. Here we describe Dyad-seq, a method that combines enzymatic detection of modified cytosines with nucleobase conversion techniques to quantify the genome-wide methylation status of cytosines at the resolution of individual CpG dinucleotides. We find that the fidelity of DNMT1-mediated maintenance methylation is directly related to the local density of DNA methylation, and for genomic regions that are lowly methylated, histone modifications can dramatically alter the maintenance methylation activity. Further, to gain deeper insights into the methylation and demethylation turnover dynamics, we extended Dyad-seq to quantify all combinations of 5mC and 5-hydroxymethylcytosine (5hmC) at individual CpG dyads to show that TET proteins preferentially hydroxymethylate only one of the two 5mC sites in a symmetrically methylated CpG dyad rather than sequentially convert both 5mC to 5hmC. To understand how cell state transitions impact DNMT1-mediated maintenance methylation, we scaled the method down and combined it with the measurement of mRNA to simultaneously quantify genome-wide methylation levels, maintenance methylation fidelity and the transcriptome from the same cell (scDyad&T-seq). Applying scDyad&T-seq to mouse embryonic stem cells transitioning from serum to 2i conditions, we observe dramatic and heterogenous demethylation and the emergence of transcriptionally distinct subpopulations that are closely linked to the cell-to-cell variability in loss of DNMT1-mediated maintenance methylation activity, with regions of the genome that escape 5mC reprogramming retaining high levels of maintenance methylation fidelity. Overall, our results demonstrate that while distinct cell states can substantially impact the genome-wide activity of the DNA methylation maintenance machinery, locally there exists an intrinsic relationship between DNA methylation density, histone modifications and DNMT1-mediated maintenance methylation fidelity that is independent of cell state.

## Introduction

Inheritance of the epigenetic mark DNA methylation (5-methylcytosine or 5mC) during cell division is critical to ensure that cellular identity is transmitted from mother to daughter cells. While inheritance of DNA methylation is primarily performed by the maintenance DNA methyltransferase 1 (DNMT1) protein by copying methylated cytosines in a CpG sequence context (5mCpG) from the old to new DNA strand, recent work has suggested that DNMT1 displays imprecise maintenance activity^1–3^. However, it remains unclear if the fidelity of DNMT1 varies at different genomic regions as well as when cells transition from one state to another. For example, one of the most dramatic illustrations of this differential DNMT1 activity is the genome-wide erasure of 5mC while methylation is maintained at imprinted loci during mammalian preimplantation embryogenesis^3^. Further, we recently showed that the genome-wide loss of 5mC during mouse and human preimplantation development is heterogenous, with a subset of cells experiencing global passive demethylation, where 5mC is not faithfully copied to the newly synthesize DNA strands^4^. Therefore, more generally, it remains unknown how cell-to-cell heterogeneity in DNMT1 maintenance methylation activity impacts cellular phenotypes. Investigating these questions remain a challenge due to a lack of experimental techniques. While the methylation status of CpG dinucleotides, a readout of DNMT1 maintenance activity, can be investigated using hairpin-bisulfite sequencing or extensions of this method, where complimentary DNA strands are physically linked, these techniques typically have low efficiency and are challenging to scale down to a single-cell resolution^5–7^. Further, physically linking the two opposing strands using a hairpin prevents direct investigation of 5mC on one strand and the oxidized derivative 5-hydroxymethylcytosine (5hmC) on the other strand of a single DNA molecule. Therefore, to investigate how the chromatin landscape and cellular state tune DNMT1 maintenance methylation fidelity and DNA methylation dynamics, we have developed a new technology (Dyad-seq) that integrates enzymatic detection of modified cytosines with traditional nucleobase conversion techniques to quantify all combinations of 5mC and 5hmC at individual CpG dyads. Finally, we have scaled down Dyad-seq and integrated it with simultaneous quantification of the transcriptome from the same cell to gain deeper insights into how DNA methylation and DNMT1-mediated maintenance methylation regulates gene expression.

## Results

### Dyad-seq variants can quantify all combinations of 5mC and 5hmC on a genome-wide scale at the resolution of individual CpG dyads

To address the above questions, we developed four different versions of Dyad-seq in bulk, two where 5mC is selected on one strand through the digestion of DNA with MspJI followed by the interrogation of 5mC or 5hmC on the CpG site of the opposing strand (M-M-Dyad-seq and M-H-Dyad-seq, respectively), and two where 5hmC is selected on one strand through the digestion of DNA with AbaSI followed by the interrogation of 5mC or 5hmC on the CpG site of the opposing strand (H-M-Dyad-seq and H-H-Dyad-seq, respectively)^4, 8^ (Fig. 1, 2a and Extended Data Fig. 1). Briefly, after digestion of DNA with either MspJI or AbaSI, the bottom strand of the fragmented molecules are captured by ligation to a double-stranded adapter containing the corresponding overhang, a sample barcode, a unique molecule identifier (UMI), and a PCR amplification sequence. Next, to detect unmodified or methylated cytosines on the opposing DNA strand, samples are either treated enzymatically with Tet2, T4-phage beta-glucosyltransferase (T4-BGT) and APOBEC3A or with sodium bisulfite to convert unmodified cytosines to uracils while methylated cytosines remain unchanged (M-M-Dyad-seq and H-M-Dyad-seq)^9^. While the methylated cytosines detected on the bottom strand in these two methods includes both 5mC and 5hmC, similar to other bisulfite-based sequencing methods, as the percentage of methylated cytosines that are 5hmC in most cell types is extremely small, it is a reasonable approximation to call these methylated cytosines as 5mC. Further, slight modification to the enzymatic conversion reaction by adding T4-BGT and APOBEC3A only results in the detection of 5hmC on the opposing DNA strand (M-H-Dyad-seq and H-H-Dyad-seq)^10–14^. The bottom strand of the adapter is unaffected by these conversion reactions as it is devoid of cytosines^15^. Next, extension using a random nonamer primer is used to incorporate part of the Illumina read 2 adapter sequence, and the resulting molecules are then PCR amplified and sequenced on an Illumina platform. From the sequencing data, the location of the methylated or hydroxymethylated cytosine on the non-amplified strand, detected by the endonuclease MspJI or AbaSI, can be inferred based on its distance from the adapter, while the methylation/hydroxymethylation status of the opposing CpG site, as well as other cytosines on this strand, can be determined directly from the sequencing results of the conversion reaction (Fig. 1 and Extended Data Fig. 1). Thus, Dyad-seq not only enables measurement of the percentage of 5mC or 5hmC at a single-base resolution, similar to that obtained from bisulfite sequencing-based approaches, but also enables quantification of the percentage of 5mC or 5hmC maintenance at individual CpG dyads. Finally, M-H-Dyad-seq and H-M-Dyad-seq allow for the direct detection of two different epigenetic marks at individual CpG dyads, measurements that are not possible with hairpin bisulfite-based techniques.

**Figure 1:**
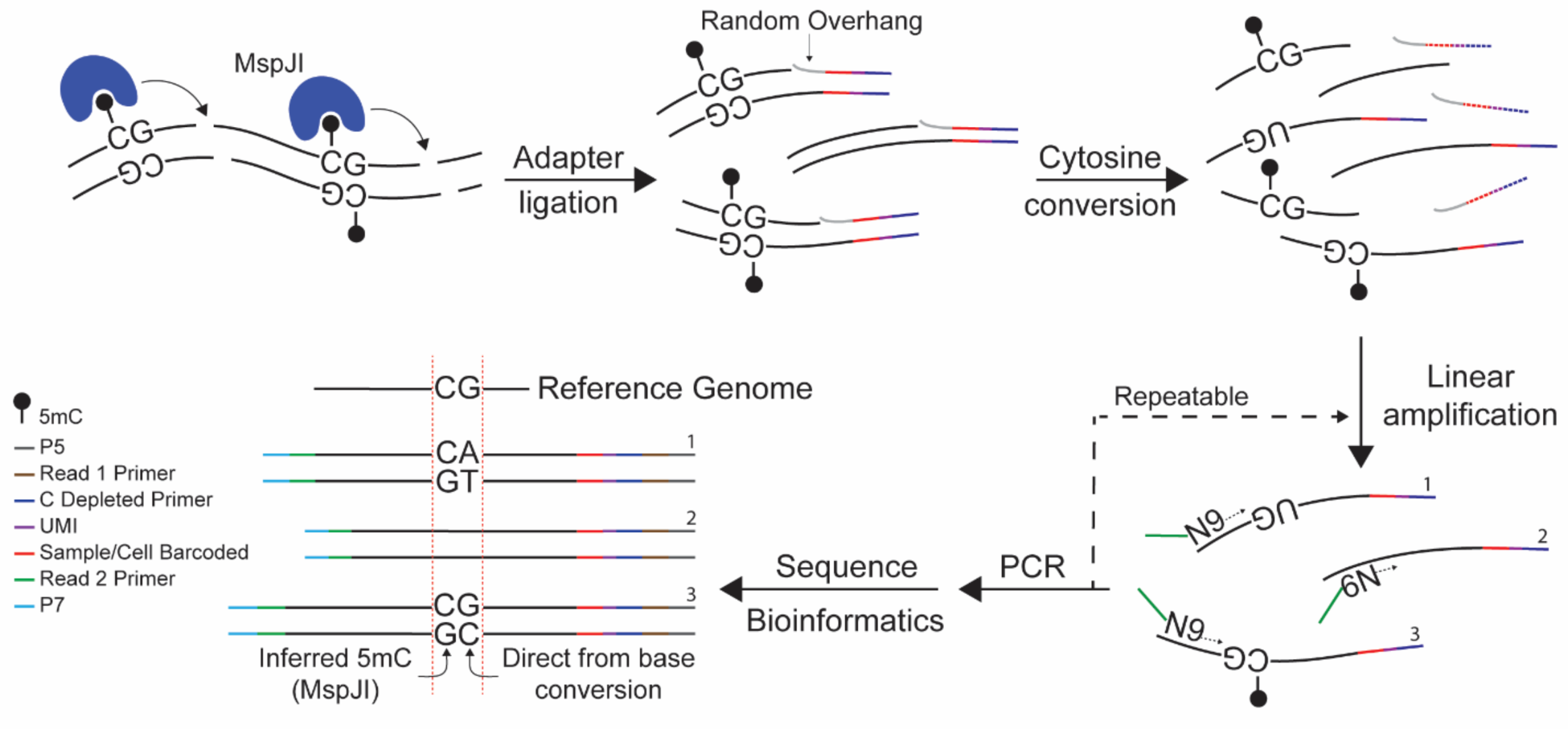
Schematic of M-M-Dyad-seq. In M-M-Dyad-seq, the schematic shows that methylated cytosines at CpG dyads are detected using MspJI digestion followed by nucleobase conversion to interrogate the methylation status of the cytosine on the opposing strand of the dyad.

To validate M-M-Dyad-seq, we first compared mouse embryonic stem cells (mESC) grown with or without Decitabine, a cytosine analog known to directly inhibit DNMT1 activity^16^. Treatment with Decitabine for 24 hours resulted in a global loss of DNA methylation as well as a dramatic reduction in 5mCpG maintenance, quantified as the fraction of CpG sites that are symmetrically methylated, demonstrating that M-M-Dyad-seq can be used to measure genome-wide DNA methylation levels and the fidelity of DNMT1-mediated maintenance methylation (Extended Data Fig. 2a,b). In addition, CpHpG maintenance methylation was very low in both conditions, consistent with the known preference of DNMT1 to maintain methylation only at CpG sites in mammalian cells (Extended Data Fig. 2c)^3^. Further, comparison to hairpin-bisulfite sequencing showed that M-M-Dyad-seq showed similar maintenance methylation levels on a genome-wide scale (Extended Data Fig. 2d)^6^. Furthermore, the genome-wide methylation levels quantified by M-M-Dyad-seq showed strong monotonic correlation with bulk bisulfite sequencing (Pearson *r* = 0.94, Spearman ! = 0.94) (Extended Data Fig. 2e)^17^. Finally, the distribution of DNA methylation over different genomic elements in M-M-Dyad-seq was broadly similar to that observed in whole-genome bisulfite sequencing as well as in scMspJI-seq, a strand-specific method that we recently developed for quantifying DNA methylation (Extended Data Fig. 2g-j)^4, 18^. Overall, these results suggested that M-M-Dyad-seq can be used to quantify DNA methylation and DNMT1-mediated maintenance methylation levels on a genome-wide scale.

After validating the technique, we next applied Dyad-seq to an *in vitro* model of epigenetic reprogramming by transitioning mESCs cultured in serum containing media supplemented with leukemia inhibitory factor (LIF) (denoted by ‘SL’)) to a serum-free media (basal media) containing LIF and two inhibitors, GSK3i (CHIR99021) and MEKi (PD0325901) (denoted by ‘2iL’)) (Fig. 2b-g)^19^. These two states of mESCs are interconvertible, with SL mESCs displaying high levels of methylation that become hypomethylated in 2iL^17^. After transitioning SL mESCs to 2iL for 48 hours, we similarly observed a global loss of 5mCpG that is accompanied with a corresponding reduction in DNMT1-mediated maintenance methylation activity (Fig. 2b,f). Similarly, when we bin the genome into 100 kb regions, we observed that in greater than 95% of the bins, reduced 5mCpG in 2iL conditions is associated with reduced maintenance methylation, consistent with previous observations that passive demethylation is the main contributor to this demethylation process (Extended Data Fig. 3a)^20–22^. To further investigate the role of different modes of demethylation during this global erasure of the methylome, we transitioned SL mESCs to different media conditions for 48 hours and performed all four variants of Dyad-seq (Fig. 2b-g and Extended Data Fig. 3b,c). In the basal media containing neither of the two inhibitors or LIF (denoted by ‘No’), cells spontaneous differentiated with a rapid increase in both the absolute levels of 5mCpG as well as DNMT1-mediated maintenance methylation (Fig. 2b,f and Extended Data Fig. 3b). In the case where LIF alone (denoted by ‘BL’) or a combination of LIF and GSK3i (denoted by ‘GL’) were added to the basal media, we observed limited changes in both the absolute levels of 5mCpG and maintenance methylation activity (Fig. 2b,f and Extended Data Fig. 3b). In contrast, basal media containing LIF and MEKi (denoted by ‘ML’) induced an even larger decrease in both the global levels of 5mCpG and DNMT1-mediated maintenance methylation fidelity than 2iL (Fig. 2b,f and Extended Data Fig. 3b). Interestingly, although the maintenance methylation fidelity decreased in both ML and 2iL conditions compared to SL and No, M-H-Dyad-seq showed that the fraction of 5mC-containing dyads that were paired with 5hmC on the opposing strand were similar for No, SL, 2iL and ML (Fig. 2c). Similarly, the genome-wide levels of 5hmC were also mostly unchanged between these different conditions (Fig. 2g and Extended Data Fig. 3c). Overall, these results highlight that the global gain or loss of methylation in this system is not dependent on TET-mediated activity but is closely linked to the fidelity of DNMT1-mediated methylation maintenance.

**Figure 2:**
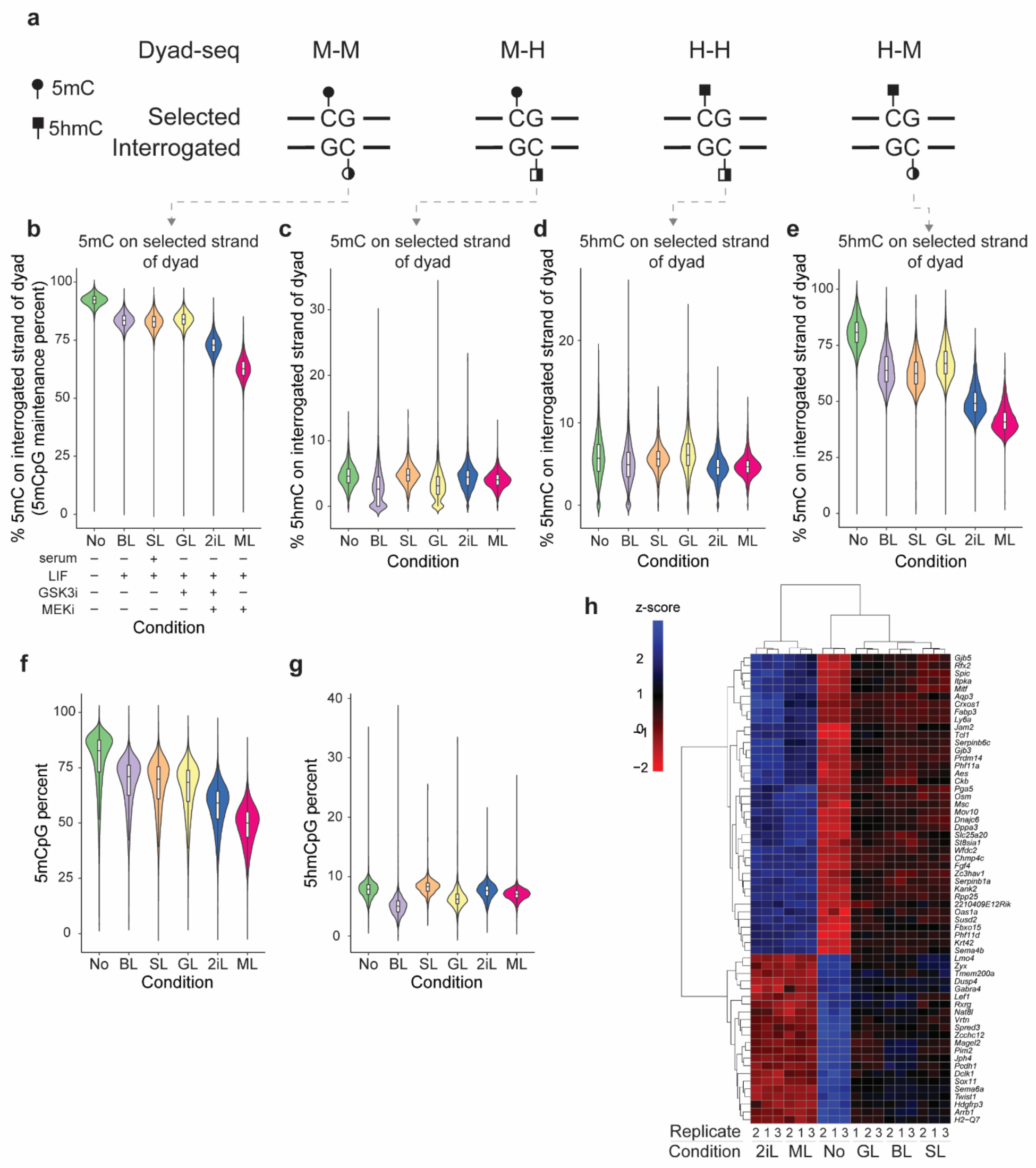
Dyad-seq variants enable genome-wide detection of all combinations of 5mC and 5hmC at individual CpG dinucleotides. (**a**) Schematic describing four different versions of Dyad-seq. M-M-Dyad-seq profiles 5mC on one strand and if the cytosine on the opposing strand of the dyad is methylated or not. M-H-Dyad-seq profiles 5mC on one strand and if the cytosine on the opposing strand of the dyad is hydroxymethylated or not. H-H-Dyad-seq profiles 5hmC on one strand and if the cytosine on the opposing strand of the dyad is hydroxymethylated or not. H-M-Dyad-seq profiles 5hmC on one strand and if the cytosine on the opposing strand of the dyad is methylated or not. (**b**) 5mCpG maintenance, quantified as the percentage of CpG dinucleotides that are symmetrically methylated, is shown for mESCs grown under different conditions. M-M-Dyad-seq is used to estimate 5mCpG maintenance. (**c**) M-H-Dyad-seq shows the percentage of 5mC that are paired with 5hmC at CpG dyads. (**d**) H-H-Dyad-seq shows the percentage of 5hmC that are paired with 5hmC at CpG dyads. (**e**) H-M-Dyad-seq shows the percentage of 5hmC that are paired with 5mC at CpG dyads. (**f**) Genome-wide 5mCpG levels quantified using M-M-Dyad-seq. (**g**) Genome-wide 5hmCpG levels quantified using M-H-Dyad-seq. In panels (b,f), the violin plots are made using 100 kb bins, and in panels (c-e,g), the violin plots are made using 1 Mb bins. In panels (b-g), mESCs are grown under SL conditions before being transitioned to the following conditions for 48 hours: No – basal media, BL – basal media with LIF, SL – serum containing media with LIF, GL – basal media with LIF and GSK3i, 2iL – basal media with LIF, GSK3i and MEKi, or ML – basal media with LIF and MEKi. (**h**) Heatmap of differentially expressed genes with a putative role in regulating DNMT1-mediated maintenance fidelity.

As we observed that the transition of SL mESCs to the No condition was associated with an increase in both 5mCpG and DNMT1-mediated maintenance methylation, while the 2iL and ML conditions were associated with a decrease in both 5mCpG and DNMT1-mediated maintenance methylation, we hypothesized that quantifying the transcriptome in these different conditions could potentially be used to identify new factors that are involved in regulating the DNA maintenance methylation machinery. For example, 2iL and ML conditions involve the inhibition of the MAPK/ERK pathway, which has previously been shown to reduce protein levels of DNMT1 in multiple systems, including during the transition of mESCs from SL to 2iL^22–24^. Therefore, we performed RNA-seq on all conditions, and as expected, found each condition to be transcriptionally distinct (Extended Data Fig. 3d,e). As DNMT1 displayed reduced maintenance methylation fidelity in the ML and 2iL conditions, but an increase in the No condition, we reasoned that putative genes involved in tuning maintenance methylation could be identified as those that are upregulated or downregulated in ML and 2iL when compared to No, but are expressed at intermediate levels in SL, GL, and BL conditions. Using this combinatorial criteria, we identified 61 differentially expressed genes, 39 of which were highly expressed in the 2iL and ML conditions with enrichment in pathways associated with pluripotency, negative cell cycle regulation, and blastocyst development, while 22 genes were highly expressed in the No condition with enrichment in pathways associated with the negative regulation of ERK1 and ERK2 cascade and mesenchymal cell differentiation (Fig. 2h and Extended Data Fig. 3f-g)^25^. Notably, our screen identified *Dppa3* (Developmental pluripotency associated 3) as one of the hits that is highly expressed in the ML and 2iL condition (Fig. 2h and Extended Data Fig. 3e). Previous studies have found that ectopic expression of DPPA3 leads to global hypomethylation, while *Dppa3* knockout leads to global hypermethylation^21, 26^. Further, DPPA3 has even been shown to directly bind the PHD domain of UHRF1 (Ubiquitin like with PHD and ring finger domains 1), a critical partner of DNMT1 necessary for 5mCpG maintenance, and displaces it from chromatin, thus inhibiting methylation maintenance^21^. The identification of a previously well-characterized factor involved in DNA methylation maintenance suggests that several of the other 60 genes identified in our screen, by combining Dyad-seq with RNA-seq, could potentially reveal novel regulators of DNA methylation maintenance fidelity and pluripotency.

We also used these experiments to further investigate the dynamics and relationship between DNA methylation and DNA hydroxymethylation at individual CpG dyads. While M-M-Dyad-seq showed as expected that a majority of 5mC existed as symmetrically methylated dyads instead of hemimethylated dyads, H-H-Dyad-seq showed that 5hmC was found to infrequently occur as symmetrically hydroxymethylated dyads (Fig. 2b,d). In contrast to this, H-M-Dyad-seq showed that 5hmC sites had high levels of 5mC on the CpG site of the opposing DNA strand, which showed similar trends to the global levels of 5mC among conditions (Fig. 2e,f and Extended Data Fig. 2f,3b). This observation is in agreement with single-molecule fluorescence resonance energy transfer experiments, which while lacking locus-specific information, globally identified that approximately 60% of 5hmC sites exist in a 5hmC/5mC dyad state in mESC^27^. These measurements strongly suggest that in mESCs TET proteins hydroxymethylate only one of the two 5mC sites in a symmetrically methylated dyad and do not sequentially convert both 5mC to 5hmC. This result is consistent with previous *in vitro* work showing that TET proteins have a stronger binding affinity for symmetrically methylated dyads over hemimethylated dyads, and a crystal structure that indicates the non-reactive cytosine is not involved in protein-DNA contacts^28, 29^. In summary, these experiments show that the combination of the four Dyad-seq variants can provide insights into DNA methylation/hydroxymethylation turnover at a single-base and genome-wide resolution that was previously not possible.

### Fidelity of DNA methylation maintenance machinery is linked to the local density of the methylome and the landscape of histone modifications

As the genome-wide methylation levels were strongly related to the overall maintenance methylation activity for mESCs grown under different conditions, we next explored how methylation levels were linked to DNMT1-mediated maintenance methylation fidelity for different genomic regions within the same cell state (Fig. 2b,f). For example, as has been shown previously, we find that compared to other genomic loci within the same cell state, intracisternal A particles (IAPs) display high 5mCpG levels in SL as well as in the globally hypomethylated 2iL and ML conditions (Fig. 3a)^20, 30^. Interestingly, these elevated 5mCpG levels at IAPs are associated with higher maintenance methylation activity, suggesting that high levels of methylation locally could be linked to increased DNMT1-mediated maintenance methylation fidelity (Fig. 3b). More generally, we find that genomic regions with increasing methylation levels are correlated with higher maintenance methylation fidelity, with a more pronounced affect in regions with higher CpG density, such as CpG islands (Fig. 3c,d and Extended Data Fig. 4a,b). Consistent with recent findings, our results from M-M-Dyad-seq suggests that there exists a tight coupling and a positive feedback loop between local methylation density and DNMT1-mediated maintenance methylation activity^2^.

**Figure 3:**
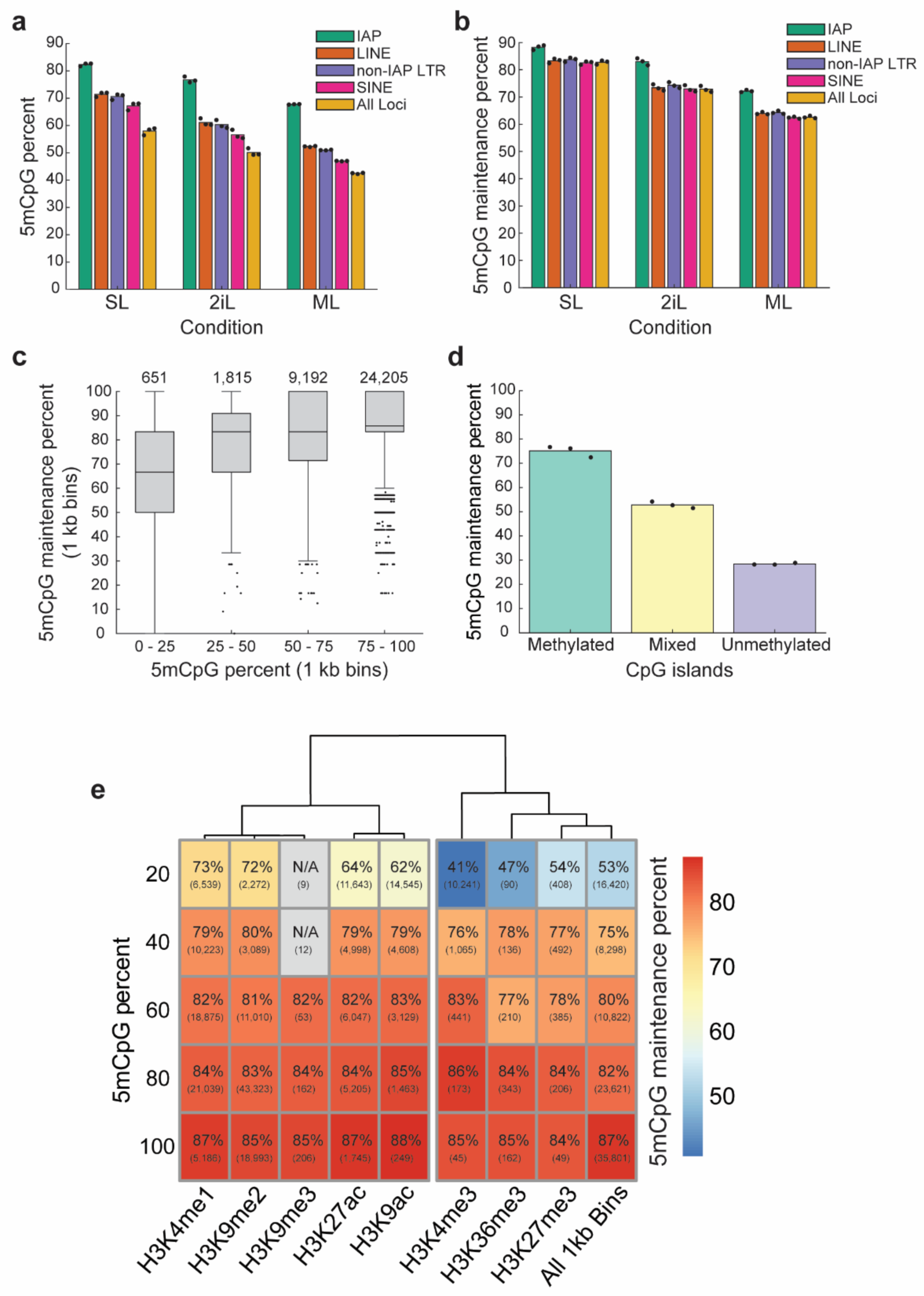
DNA methylation maintenance activity is linked to the local density of the methylome and the distribution of histone modifications. (**a**) Bar plots show 5mCpG levels estimated using M-M-Dyad-seq at various repetitive elements for mESCs after 48-hours in the indicated media conditions. Dots indicate independent biological replicates (n = 3). (**b**) Bar plots show 5mCpG maintenance fidelity estimated using M-M-Dyad-seq at various repetitive elements for mESCs grown under different conditions for 48 hours. IAPs are more resistant to demethylation that other genomic regions and show higher 5mCpG maintenance levels. Dots indicate independent biological replicates (n = 3). (**c**) Box plot of 5mCpG maintenance levels in 1 kb genomic bins over varying absolute methylation levels. 1 kb regions in which at least 5 unique CpG dyads are detected were included in this panel. The number of bins in each category is denoted above each boxplot. Data from mESCs grown in SL and profiled using M-M-Dyad-seq is shown in this panel. (**d**) 5mCpG maintenance levels at CpG islands separated based on absolute methylation levels. ‘Methylated’ indicates CpG islands with greater than 20% 5mCpG levels, ‘mixed’ indicates 5mCpG levels between 10 and 20%, and ‘unmethylated’ indicates CpG islands with less than 10% 5mCpG levels. Data from mESCs grown in SL and profiled using M-M-Dyad-seq is shown in this panel. (**e**) Heatmap of 5mCpG maintenance fidelity in SL grown mESCs at genomic regions enriched for various histone marks. Numbers within parenthesis indicate the total number of regions analyzed in the meta-region.

Based on this discovery that 5mCpG maintenance can be tuned by the methylation levels at different genomic loci, we hypothesized that the genome-wide landscape of histone modifications could also impact DNMT1-mediated maintenance methylation. We found that highly methylated genomic regions are associated with high 5mCpG maintenance, independent of the histone modification at those loci (Fig. 3e). Surprisingly, however, we discovered that for regions that are lowly methylated, the presence of a particular histone mark can dramatically alter the maintenance methylation fidelity (Fig. 3e and Extended Data Fig. 4c-k). For example, we found regions enriched for the repressive mark H3K9me2 are associated with higher maintenance methylation fidelity than a randomly selected bin at similar methylation levels (Fig. 3e and Extended Data Fig. 4d). This is consistent with previous observations that UHRF1 can specifically bind H3K9me2 with high affinity, providing a mechanistic rationale for the recruitment of DNMT1 and higher maintenance seen in these regions^31–33^. Interestingly, we identify that enhancers marked by H3K4me1 or H3K27ac, and active promoters/enhancers marked by H3K9ac also have increased DNMT1-mediated maintenance methylation fidelity (Fig. 3e and Extended Data Fig. 4e,h,k,l). In contrast, H3K4me3, a mark found at active gene promoters, had lower levels of 5mCpG maintenance than the average genome-wide maintenance (Fig. 3e and Extended Data Fig. 4j,l)^34^. Overall, these results demonstrate that for regions of the genome that are lowly methylated, histone modifications can significantly alter maintenance methylation fidelity from approximately 40% to over 70% of the dyads being symmetrically methylated.

### scDyad&T-seq enables joint profiling of the methylome, maintenance methylation fidelity and the transcriptome from the same cell

In addition to the dramatic heterogeneity we observed in maintenance methylation across different genomic contexts, we next wanted to understand how cell-to-cell variability in DNMT1-mediated maintenance methylation fidelity could impact cellular phenotypes within a population. Addressing this question has been challenging as current hairpin bisulfite techniques have not successfully been applied to single cells. To overcome this limitation, we scaled down M-M-Dyad-seq to a single-cell resolution (termed as scDyad-seq). As proof-of-concept, cells treated with Decitabine for 24 hours experienced a global loss of DNA methylation and a dramatic reduction in the fraction of CpG dyads that were symmetrically methylated, while showing no changes in 5mCpHpG maintenance, as expected (Extended Data Fig. 5a,b). Further, to directly link heterogeneity in DNA methylation and DNMT1-mediated maintenance methylation fidelity to gene expression variability, we extended the method to simultaneously also capture the transcriptome (scDyad&T-seq) using an amplification strategy we recently developed ^35^. Briefly, single cells are sorted into 384-well plates and the mRNA is converted to cDNA, as described previously, using poly-A primers with overhangs that contain a cell-specific barcode, UMI, 5’ Illumina adapter and T7 promoter (Fig. 4a)^35^. Following this, the next steps are similar to M-M-Dyad-seq based digestion and ligation where the genomic DNA within the reaction mix is glucosylated, digested with the restriction enzyme MspJI and ligated to Dyad-seq double-stranded adapters that contain a cell-specific barcode, UMI and cytosine-depleted PCR handle. Thereafter, as the genomic DNA and cDNA within individual cells are uniquely tagged, the samples are pooled and subjected to *in vitro* transcription (IVT). IVT only amplifies T7 promoter containing cDNA molecules, and these amplified RNA molecules are pulled down using biotinylated poly-A oligonucleotides and magnetic streptavidin beads, as described previously^35^. The amplified RNA molecules are then used to prepare Illumina libraries, enabling quantification of the transcriptome of single cells^35, 36^. The flowthrough from the bead separation contains unamplified Dyad-seq adapter ligated genomic DNA molecules that are then subjected to nucleobase conversion, random linear amplification, and PCR, similar to that described above for M-M-Dyad-seq, to quantify the methylome and the methylation status of CpG dyads on a genome-wide scale from the same single cells (Fig. 4a).

**Figure 4:**
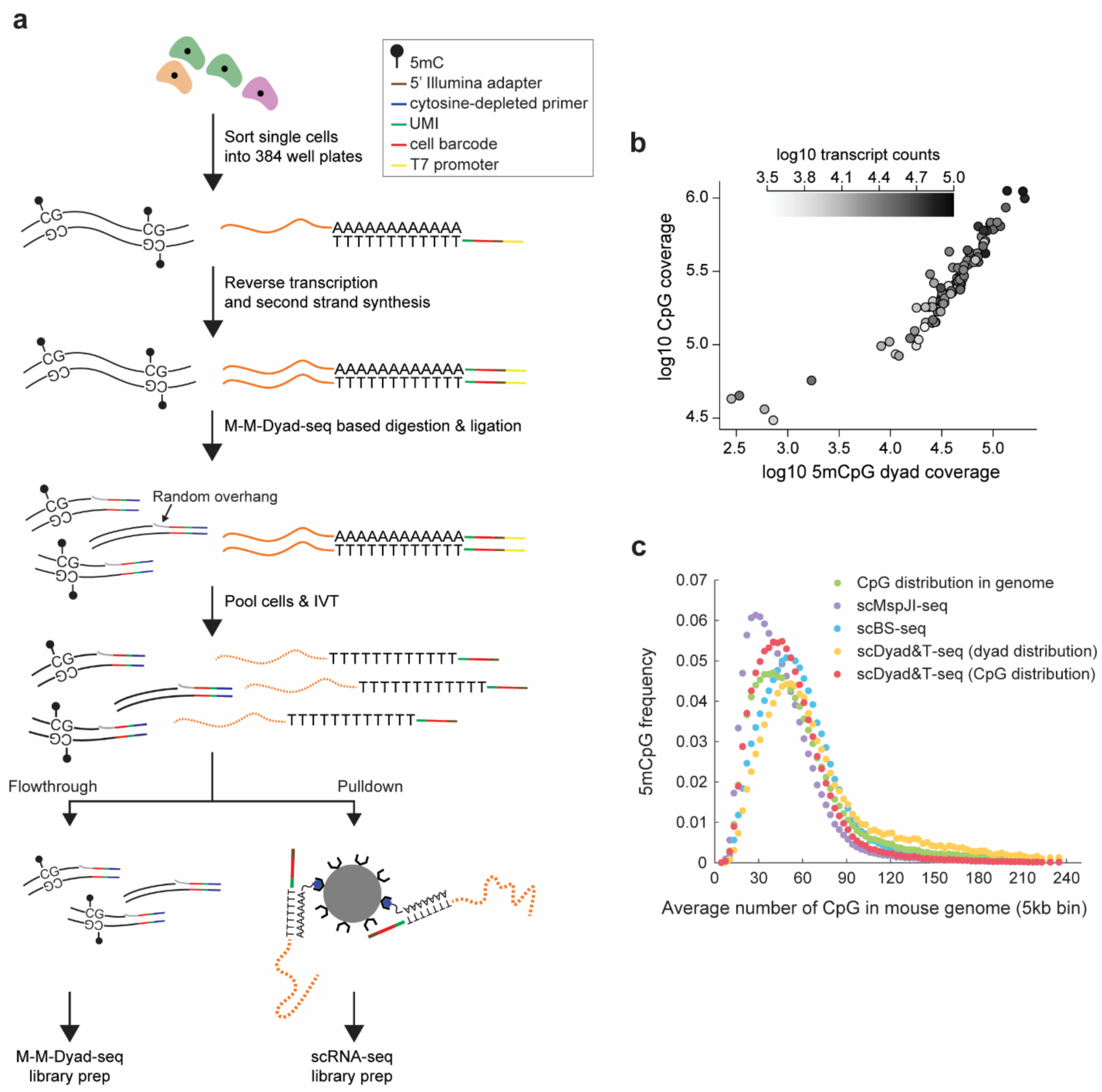
scDyad&T-seq enables joint profiling of the methylome, maintenance methylation fidelity and the transcriptome from the same cell. (**a**) Schematic of scDyad&T-seq. By scaling down M-M-Dyad-seq and combining it with the capture of mRNA, scDyad&T-seq can simultaneously quantify DNA methylation levels, maintenance methylation levels and the transcriptome from the same cell in a high-throughput format. Black, solid orange and dotted orange lines correspond to genomic DNA, mRNA and amplified RNA, respectively. (**b**) Panel shows the coverage of CpG dinucleotides that provide information on maintenance methylation (5mCpG dyad coverage), and coverage of CpG sites that enable quantification of absolute methylation levels (CpG coverage), together with the number of unique transcripts detected in individual cells. Note that 5mCpG dyad coverage and CpG coverage are derived from non-overlapping regions of the same sequencing read; therefore, the total number of CpG sites interrogated in a cell can be obtained by summing of 5mCpG dyad coverage and CpG coverage. (**c**) Distribution of 5mCpG sites that are detected by different methods (scDyad&T-seq, scMspJI-seq and scBS-seq) over genomic regions of varying CpG density^4, 18^. The green curve shows the distribution of CpG sites over 5 kb bins in the mouse genome.

We first applied scDyad&T-seq to serum grown mESCs cells to detect up to 75,835 unique transcripts per cell, and the methylation status of up to 1,118,393 CpG sites per cell, together with the additional detection of the maintenance methylation status of up to 203,620 CpG dyads per cells (with an average of 25,066 unique transcripts per cell (5,825 genes/cell), covering the methylation status of 328,967 CpG sites on average per cell and the maintenance methylation status of an additional 51,650 CpG dyads on average per cell) (Fig. 4b). In comparison, while scM&T-seq, a method that captures the methylome and transcriptome from the same cell, covered more CpG sites and genes per cell, these libraries were proportionally sequenced at a much higher depth than the scDyad&T-seq libraries (Extended Data Fig. 5c)^37^. Further, as scM&T-seq requires physical separation of genomic DNA and mRNA prior to barcoding, amplification and library preparation, this method is more challenging to scale up to sequence thousands of cells per day, and unlike scDyad&T-seq cannot capture the methylation status of individual CpG dyads in single cells. Finally, the distribution of CpG sites covered by scDyad&T-seq was similar to their distribution in the genome and that detected by single-cell bisulfite sequencing, suggesting that scDyad&T-seq can capture the genome-wide methylome and maintenance methylation landscape in single cells (Fig. 4c)^18^.

We further compared scDyad&T-seq to scMspJI-seq, a method we recently developed for strand-specific quantification of 5mC^4^. While scMspJI-seq does not have the resolution of individual CpG dyads, it can be used to estimate the extent of asymmetry in DNA methylation between two strands of DNA over a large genomic region^4^. We previously quantified this strand-specific DNA methylation using a metric known as strand bias, defined as the number of methylated cytosines on the plus strand divided by the total number of methylated cytosines on both DNA strands, with deviations from a score of 0.5 indicating asymmetric DNA methylation between the two strands of DNA^4^. Therefore, we directly compared the individual-CpG-dyad (or 5mCpG maintenance) resolution afforded by scDyad&T-seq to the strand bias score that can be obtained from both scDyad&T-seq as well as scMspJI-seq (as both these methods employ the same restriction enzyme MspJI). Overall, we found agreement between both measurements, with high 5mCpG maintenance associated with no strand bias (Fig. 5a,b and Extended Data Fig. 6a). Notably, however, we found that the strand bias score in general can only identify cells that experience high levels of strand-specific asymmetry in DNA methylation and, unlike the individual-CpG-dyad resolution of scDyad&T-seq, is not able to capture the wide gradient in strand-specific DNA methylation in single cells (Fig. 5a). Further, this limited sensitivity when using the strand bias score in scMspJI-seq possibly also arises from a lack of allele-specific resolution of this technique. For example, while low levels of 5mCpG maintenance (Mean 5mCpG maintenance = 25.4%) in cell P7L3.69 was associated with strand bias scores that substantially deviated from 0.5 as expected, cell P7L4.78 displaying similarly low levels of 5mCpG maintenance (Mean 5mCpG maintenance = 30.7%) showed very little strand bias (Fig. 5b). Similarly, we next used scDyad&T-seq to test the performance of a computational method for estimating maintenance methylation from nucleobase conversion-based 5mC sequencing data^38^. While the computational metric performed reasonably well for cells that displayed low 5mCpG maintenance, its accuracy was lower for cells with high levels of 5mCpG maintenance (Spearman correlation, ! = 0.42) (Extended Data Fig. 6b). Together these results demonstrate that scDyad&T-seq is a highly sensitive method for assessing the maintenance methylation status at individual CpG dyads in single cells, and when combined with the readout of absolute levels of methylation and the transcriptome from the same cell, this method could potentially allow us to directly link DNA methylation dynamics and heterogeneity to cellular phenotypes.

**Figure 5:**
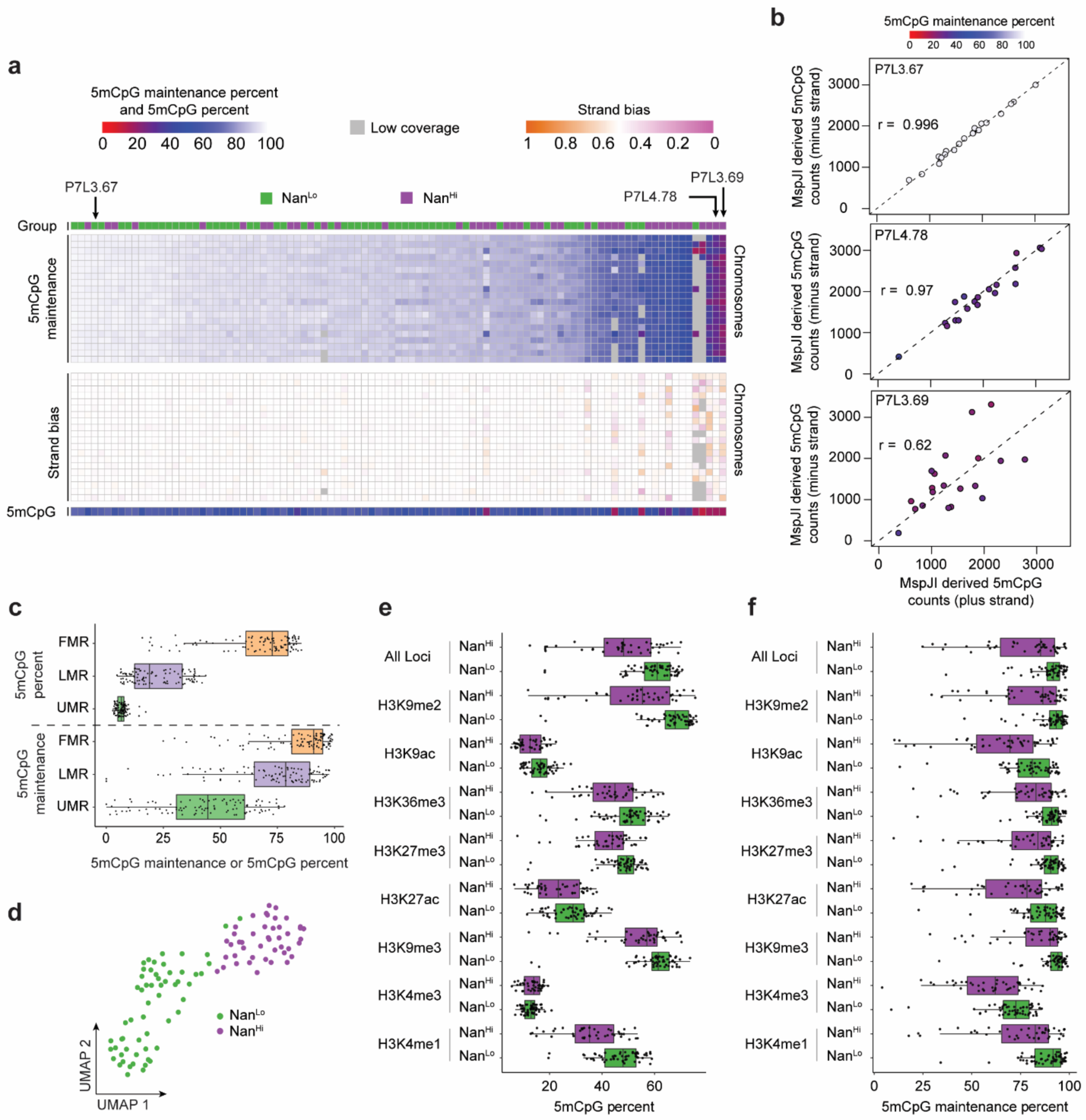
scDyad&T-seq can directly correlate heterogeneity in the methylome and DNMT1-mediated maintenance fidelity to transcriptional variability in single cells. (**a**) Heatmap comparing 5mCpG maintenance over individual chromosomes in mESCs computed using scDyad&T-seq with the strand bias metric that can be estimated from techniques such as scMspJI-seq from the same single cells. The heatmap shows that the 5mCpG maintenance estimated from scDyad&T-seq displays increased sensitivity in quantifying strand-specific DNA methylation compared to the strand bias metric obtained from scMspJI-seq^4^. The transcriptional group individual cells belong to (top) and their genome-wide 5mCpG methylation levels (bottom) are also reported in this panel. (**b**) Example of strand-specific methylation of three cells, P7L3.67, P7L4.78 and P7L3.69, from panel (a). For cell P7L3.67, similar levels of 5mCpG detected on the plus and minus strand of each chromosome by the enzyme MspJI is in agreement with the high levels of 5mCpG maintenance quantified using scDyad&T-seq. In contrast, cells P7L4.78 and P7L3.69 show very similar levels of 5mCpG maintenance computed using scDyad&T-seq but display substantial differences when MspJI-based quantification is used to estimate strand-specific methylation, indicating the limited accuracy of the strand bias score used in scMspJI-seq. A low Pearson’s correlation indicates deviations from a strand bias score of 0.5. Color of the data points indicates 5mCpG maintenance percent of individual chromosomes estimated using scDyad&T-seq. (**c**) DNA methylation and 5mCpG maintenance levels at different genomic regions stratified by Stadler *et al*. as fully methylated regions (FMR), lowly methylated regions (LMR), and unmethylated regions (UMR)^52^. Data points represent individual cells. (**d**) UMAP visualization of serum grown mESCs based on the single-cell transcriptomes obtained from scDyad&T-seq. (**e**) 5mCpG levels in regions marked by specific histone modifications. (**f**) 5mCpG maintenance levels in regions marked by specific histone modifications. In panels (e,f), cells are grouped based on the transcriptional clusters shown in panel (d).

### Distinct cell states display varied maintenance methylation fidelity and genome-wide DNA methylation landscapes

After establishing that scDyad&T-seq can accurately quantify maintenance methylation fidelity, in addition to DNA methylation and the transcriptome in single cells, we next attempted to estimate the cell-to-cell heterogeneity in the methylome and its relationship to transcriptional variability and cellular phenotypes. Surprisingly, even when considering regions of similar 5mC content in the genome based on previous bulk measurements, we observed substantial heterogeneity in both the methylation levels and 5mCpG maintenance fidelity between individual cells (Fig. 5c). Furthermore, we also noted that the relationship of higher 5mCpG maintenance fidelity at regions of the genome with higher methylation density is preserved at the single cell level (Fig. 3c-e, 5c and Extended Data Fig. 4a,b). To systematically correlate heterogeneity in the epigenome to gene expression variability and the occurrence of distinct cell states, we first used the transcriptome to identify two subpopulations in the serum grown mESCs – one high in NANOG, REX1, and ESRRB (referred to as NANOG high or ‘Nan^Hi^’) and one low in the expression of these genes (referred to as NANOG low or ‘Nan^Lo^’) (Fig. 5d, Extended Data Fig. 6c, and Supplementary Table 1). While these two well-established subpopulations in serum grown mESCs are known to be transcriptionally heterogenous with bimodal expression of key pluripotency genes, how these cell states are linked to the methylome and DNMT1-mediated maintenance methylation fidelity remains less well studied^37, 39^. We found that cells in the Nan^Hi^ state are globally hypomethylated with reduced genome-wide 5mCpG maintenance compared to the Nan^Lo^ cell state (Mann-Whitney U test, “= 8×10^−6^ and” = 3.6×10^−4^, respectively), suggesting that DNMT1-mediated maintenance methylation fidelity could play an important role in tuning the cell state in serum grown mESCs (Fig. 5e,f).

We next explored how histone modifications impact 5mCpG maintenance in the context of different cell states. Consistent with bulk findings, H3K9me2/3, H3K4me1, H3K27ac, and H3K9ac enriched genomic regions showed higher maintenance methylation fidelity while H3K4me3 marked regions showed reduced maintenance methylation fidelity (Fig. 5e,f and Extended Data Fig. 6d-g). Interestingly though, across all the histone modifications investigated, cells in the Nan^Hi^ state consistently showed reduced methylation levels and correspondingly lower 5mCpG maintenance compared to Nan^Lo^ cells, suggesting an intrinsic and mechanistic relationship between DNA methylation density and DNMT1-mediated maintenance methylation fidelity that is independent of cell state. Finally, these results also demonstrate that the cell state could play a key role in impacting the DNA methylation maintenance machinery and the resulting genome-wide methylation landscape (Fig. 5e,f).

### SL to 2iL transition in mESCs results in heterogeneous maintenance methylation activity and demethylation that is associated with the emergence of transcriptionally distinct subpopulations

To further investigate how maintenance methylation fidelity is influenced as cells transition from one state to another, we applied scDyad&T-seq to mESCs that were switched from serum containing media (SL) to 2iL media for 3, 6, or 10 days (2iL-D3, 2iL-D6, and 2iL-D10, respectively). While transition from SL to 2iL resulted in dramatic erasure of DNA methylation and loss of maintenance methylation activity, we surprisingly also found that the epigenetic reprogramming to the 2iL state is highly heterogenous with a fraction of cells still retaining high levels of methylation and 5mCpG maintenance even after 10 days in 2iL (Fig. 6a-c). Using hierarchical clustering, cells were classified as either highly or lowly methylated (mC^Hi^ and mC^Lo^), and highly or lowly maintained (Mnt^Hi^ and Mnt^Lo^), leading to 4 distinct categories of epigenetic states (Extended Data Fig. 7a,b). Superimposing the time-course data on these epigenetic states show that cells generally start off in a highly methylated and highly maintained state, with passive demethylation thereafter resulting in the loss of 5mC till they reach a lowly methylated and lowly maintained state. Surprisingly, a fraction of cells subsequently move towards a lowly methylated but highly maintained state to establish a globally hypomethylated genomic landscape that is maintained at high fidelity (Fig 6c,d). Consistent with this result, we found that regions previously identified as retaining high methylation in the 2iL state also showed higher methylation in our data, and correspondingly displayed higher maintenance methylation fidelity at all timepoints, including in SL grown cells^17^ (Fig. 6e,f). These results again highlight the intrinsic relationship between the local methylation density and maintenance methylation fidelity that is conserved through transitions in cell states (Fig. 3e, 5c,e,f and 6e,f). Further, these highly methylated regions were previously shown to correlate with H3K9me3 peaks, which are found at similar genomic regions for both SL and 2iL grown mESCs^17, 40^. This, together with our previous observation that H3K9me3-marked regions show high 5mCpG maintenance fidelity, suggests that the relationship between methylation density and maintenance methylation fidelity across cell states in this system could be coupled to H3K9me3, and that the global impairment to the 5mCpG maintenance machinery in 2iL grown mESCs is partially restored at H3K9me3-enriched regions (Fig. 3e and Extended Data Fig. 4i).

**Figure 6:**
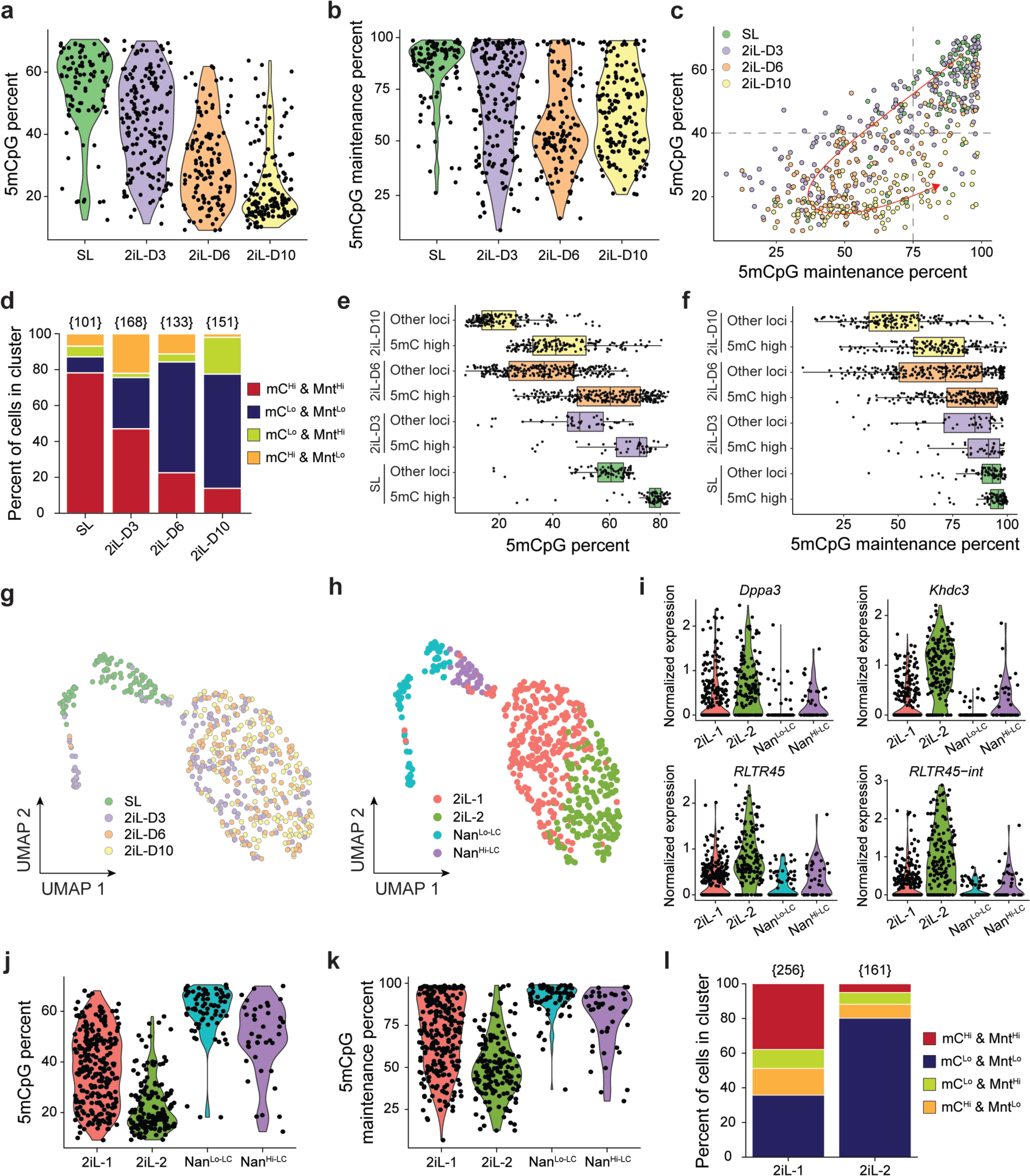
Serum to 2iL transition of mESCs involves a transient loss of methylation maintenance fidelity and is associated with the emergence of two distinct 2iL transcriptional populations. (**a,b**) Genome-wide methylation (panel a) and maintenance methylation levels (panel b) of individual mESCs cultured in SL or in 2iL conditions for 3, 6 or 10 days. (**c**) Genome-wide 5mCpG methylation and 5mCpG maintenance levels of single cells as they transition from SL to 2iL conditions. Cells transition from highly methylated and highly maintained to a lowly methylated and lowly maintained or lowly methylated and highly maintained state. (**d**) As cells transition from SL to 2iL conditions, they are classified as lowly (mC^Lo^) or highly (mC^Hi^) methylated and lowly (Mnt^Lo^) or highly (Mnt^Hi^) maintained based on hierarchical clustering, as described in Extended Data Figure 7a,b. (**e,f**) Genome-wide methylation (panel e) and maintenance methylation levels (panel f) at region previously identified by Habibi *et al*. to be highly methylated in mESCs grown in long term 2iL culture (denoted as ‘5mC high’) and at all other genomic regions (denoted as ‘Other loci’), with each point denoting a single cell^17^. (**g,h**) UMAP visualization of SL and 2iL grown cells based on the single-cell transcriptomes obtained from scDyad&T-seq, and classified by culture conditions (panel g) or by transcriptome-based clustering (panel h). (**i**) Expression levels of select genes and transposable elements, such as DPPA3, KHDC3, RLTR45, and RLTR45-int, that were found to be highly expressed in the 2iL-2 population. (**j,k**) Genome-wide methylation (panel j) and maintenance methylation levels (panel k) of single cells in different transcriptional clusters shown in panel (h). (**l**) Panel shows the percentage of 2iL-1 and 2iL-2 cells in the four groups classified based on the genome-wide methylation and maintenance methylation levels. Numbers within parenthesis indicate the total number of cells in the transcriptional clusters 2iL-1 and 2iL-2.

We next analyzed the gene expression patterns from the same cells to find that most cells in 2iL were transcriptionally similar and distinct from SL grown cells, as expected (Extended Data Fig. 7c). Further, we found that the 2iL cells did not strongly separate by the time spent in culture, suggesting that the transcriptional reprogramming is quickly activated once cells are transitioned from SL to 2iL conditions (Fig. 6g and Extended Data Fig. 7c-g)^41^. Beyond the two broad transcriptional groups of SL-like and 2iL-like cells, further clustering revealed 4 distinct populations, with two clusters transcriptionally similar to SL grown cells – Nanog low like-cells and Nanog high like-cells (referred to as ‘Nan^Lo-LC^’ and ‘Nan^Hi-LC^’, respectively) – that primarily included serum grown mESCs, and two similar to 2iL cells (referred to as ‘2iL-1’ and ‘2iL-2’) (Fig. 6h). Interestingly, we found a small group of cells from the 2iL-D3 condition cluster with the Nan^Lo-LC^ cell population. Previous work has shown that SL grown cells in the Nan^Lo^ state are less likely to survive the transition to 2iL when compared to those in the Nan^Hi^ state^42^. This low survival rate of cells in the Nan^Lo^ state could possibly arise due to an epigenetic barrier, and comparison of the 2iL-D3 cells that cluster with the SL-like cells shows that this group is characterized by high 5mCpG maintenance and correspondingly high methylation levels relative to the 2iL-D3 cells that successfully transition and cluster with the 2iL-like cells (Extended Data Fig. 8a,b). Further, consistent with recent observations, we find that the 2iL-D3 cells that cluster with the SL-like cells express low levels of *Pou5f1* (Extended Data Fig. 8c)^42^. Finally, these cells also express *Sox1*, suggesting that they begin to differentiate towards the neuroectoderm lineage, but do not survive long term culture in 2iL conditions and are not present by day 10 (Fig. 6g,h and Extended Data Fig. 8d).

In contrast, cells that successfully transition their transcriptional program to 2iL-like cells could be split into two distinct clusters, 2iL-1 and 2iL-2, with a small group of genes and transposable elements that are differentially expressed between these two sub-populations (Extended Data Fig. 8e). Notably, we found that cells in the 2iL-2 cluster expressed *Dppa3* at higher levels than the 2iL-1 cluster (Fig. 6i). Consistent with the known role of DPPA3 as an inhibitor of maintenance methylation, we found that 2iL-2 cells displayed dramatically lower DNMT1-mediated maintenance methylation fidelity, resulting in genome-wide demethylation compared to 2iL-1 cells (Fig. 6j,k)^21^. These results highlight the underlying intrinsic relationship between DNA methylation levels and maintenance methylation fidelity across the genome and how cell states preferentially occupy specific regions within the methylome density *vs.* maintenance methylation fidelity landscape (Extended Data Fig. 8f,g). Further, the globally hypomethylated state in the 2iL-2 population correlates with higher expression of endogenous retroviral elements RLTR45 and RLTR45-int, as well as higher expression of *Khdc3* (also known as *Filia*), a factor known to be involved in safeguarding genomic integrity in mESCs and preimplantation embryos^43, 44^ (Fig. 6i). Taken together, these observations suggest that reduced DNMT1-mediated maintenance fidelity in the 2iL-2 population results in genome-wide DNA demethylation, an associated loss of retroviral silencing, and genome instability. Finally, we observed that between the two 2iL subpopulations, the 2iL-2 cluster is dominated by cells in the mC^Lo^ and Mnt^Lo^ state and is weakly biased towards cells that have been cultured longer in 2iL conditions (Fig. 6l and Extended Data Fig. 8h). Similar to differences in the cell state transition potential between Nan^Lo^ and Nan^Hi^ serum grown cells, additional work in the future will be required to investigate how differences in the maintenance methylation fidelity and DNA methylation landscape between 2iL-1 and 2iL-2 subpopulations impact their differentiation and cell fate potential.

## Discussion

In summary, Dyad-seq is a generalized genome-wide approach for profiling all combinations of 5mC and 5hmC at individual CpG dyads. Using M-M-Dyad-seq, we discovered that DNMT1-mediated maintenance methylation fidelity is directly tied to local methylation levels, and for regions of the genome that have low methylation, specific histone marks can significantly modulate the maintenance methylation activity. Further, when combined with RNA-seq, we identified well-characterized factors, such as DPPA3, as well as other putative factors that are potentially involved in regulating the maintenance methylation fidelity of DNMT1. Similarly, using H-M-Dyad-seq and H-H-Dyad-seq, we found that 5hmCs are more commonly duplexed with 5mC in mESCs. These results highlight that variants of Dyad-seq, such as H-M-Dyad-seq and M-H-Dyad-seq, enable measurements that are currently not possible with hairpin bisulfite techniques, thereby providing deeper insights into the regulation of methylation and demethylation pathways in mammalian systems. To understand how changes in maintenance methylation fidelity during cell state transitions reprogram the methylome and transcriptome, we developed scDyad&T-seq to simultaneously quantify genome-wide methylation levels, maintenance methylation and mRNA from the same cells. Applying scDyad&T-seq to mESCs transitioning from SL to 2iL, we found that the epigenetic reprogramming was highly heterogenous, with the emergence of four distinct cell populations that dramatically differ in 5mCpG maintenance, DNA methylation levels, as well as gene expression. These results showed that in addition to cell identity, DNA methylation levels and histone modifications are closely tied to how faithfully the DNA methylation maintenance machinery copies methylated cytosines from one cell generation to another. Overall, scDyad-seq is an enhancement over both scMspJI-seq and single-cell bisulfite sequencing techniques, enabling high-resolution quantification of both genome-wide 5mC levels and maintenance methylation in thousands of single-cells, and when extended to scDyad&T-seq, the method can also be used to simultaneously obtain the transcriptome from the same cells (Fig. 4a,b, 5a,b, and Extended Data Fig. 6a,b, 8i,j).

### Methods Cell culture

All cells were maintained in incubators at 37°C and 5% CO_2_. Mouse embryonic stem cell line ES-E14TG2a (E14) was cultured on gelatin (Millipore Sigma, ES-006-B) coated tissue culture plates with media containing high glucose DMEM (Gibco, 10569044), 1% non-essential amino acid (Gibco, 11140050), 1% Glutamax (Gibco, 35050061), 1x Penicillin-Streptomycin (Gibco, 15140122), and 15% stem cell qualified serum (Millipore Sigma, ES-009-B). The media was frozen in aliquots and used thereafter for a maximum of 2 weeks after thawing while storing it at 4°C. Once thawed, 1 µL of beta-mercaptoethanol (Gibco, 21985023), and 1 µL of LIF (Gibco, A35933) were added for every 1 mL of thawed media. Cells were washed with 1x DPBS (Gibco, 14190250) and the media was changed daily. Cells were routinely passaged 1:6 once they reached 75% confluency using 0.25% trypsin-EDTA (Gibco, 25200056). E14 cells grown under these conditions also describe the SL experimental group. For FACS sorting, a single-cell suspension was made using 0.25% trypsin-EDTA. The trypsin was then inactivated using serum containing medium. Afterwards, the cells were washed with 1x DPBS before being passed through a cell strainer and sorted for single cells into 384-well plates.

K562 cells were grown in RPMI (Gibco, 61870036) with 10% serum (Gibco, 10437028) and 1x Penicillin-Streptomycin. When the culture reached a density of approximately 1 million cells per mL, they were split and resuspended at a density of 200,000 cells per mL. Cells were washed and FACS sorted as described for E14 cells.

### Decitabine treatment

E14 mouse embryonic stem cells were cultured as described above. Upon passage of the E14 cells, SL media was supplemented with 0.05 µM of Decitabine. After 24 hours, cells were harvested using 0.25% trypsin-EDTA. The trypsin was then inactivated using serum containing medium. The cells were washed with 1x DPBS and then resuspended in 200 µL of DPBS. Genomic DNA was extracted using the DNeasy kit (Qiagen, 69504) according to the manufacturer’s recommendations.

K562 cells were cultured as described above. Upon passage, the media was supplemented with 0.6 µM of Decitabine or DMSO (as a control). After 24 hours the cells were washed and single-cell FACS sorting was performed as described above.

### Transition from SL to 2iL and other media conditions

E14 mouse embryonic stem cells were cultured in SL conditions as described above. Upon passage, cells were resuspended in the following media depending on the condition studied. Commercial 2i media containing LIF (Millipore, SF016-200) was used for BL, GL, 2iL, and ML experiments. For 2iL, all components were used according to the manufacturer’s recommendations. For GL and ML conditions, only the GSK3B inhibitor or MEK1/2 inhibitor was added, respectively. For the BL condition, no inhibitors were added. For the No condition, commercial 2i media without LIF (Millipore, SF002-100) was used with no inhibitors added. After 24 hours, the cells were washed with 1x DPBS and the media was changed. 48 hours after the initial media switch, the cells were collected using 0.25% trypsin-EDTA, quenched using serum containing media, washed in 1x DPBS and finally resuspended in 1x DPBS. The sample was then split in half. One half was resuspended in 200 µL of DPBS for genomic DNA extraction, as described above. The other half was resuspended in 500 µL of TRIzol reagent (Invitrogen, 15596018) and total RNA was extracted according to the manufacturer’s recommendations. Experiments for each condition were performed in triplicate.

### Dyad-seq Adapters

The double-stranded Dyad-seq adapters are designed to be devoid of cytosines on the bottom strand. They contain a PCR sequence, a 4-base pair UMI, and a 10-base pair cell-specific barcode. For Dyad-seq variants that use MspJI as a restriction enzyme (M-M-Dyad-seq and M-H-Dyad-seq), the adapters contain a random 4 base pair 5’ overhang and their general design is shown below.

Top oligo: 5’-NNNN [10 bp barcode] HHWHCCAAACCCACTACACC −3’

Bottom oligo: 5’-GGTGTAGTGGGTTTGGDWDD [10 bp barcode] −3’

The sequences of the 10 bp cell-specific barcode for scDyad&T-seq are provided in Supplementary Table 2. For bulk Dyad-seq, cell-specific barcodes were used as sample-specific barcodes.

For the M-M-Dyad-seq experiments described in the manuscript, a prototype of the above design was used consisting of a 3 bp UMI and an 8 bp sample-specific barcode (Supplementary Table 3).

Top oligo: 5’-NNNN [8 bp barcode] HHHCCAAACCCACTACACC −3’

Bottom oligo: 5’-GGTGTAGTGGGTTTGGDDD [8 bp barcode] −3’

The sequences of the 8 bp sample-specific barcode for M-M-Dyad-seq are provided in Supplementary Table 3.

For Dyad-seq variants that use AbaSI as a restriction enzyme (H-H-Dyad-seq and H-M-Dyad-seq), the adapters contain a random 2 base pair 3’ overhang as shown below:

Top oligo: 5’-[10 bp barcode] HHWHCCAAACCCACTACACC −3’

Bottom oligo: 5’-GGTGTAGTGGGTTTGGDWDD [10 bp barcode] NN −3’

The sequences of the 10 bp sample-specific barcode used for H-M-Dyad-seq and H-H-Dyad-seq are provided in Supplementary Table 4.

Note: All adapters described above are unphosphorylated. The protocol for annealing the top and bottom strands to generate double-stranded adapters is described previously in scAba-seq^8^.

### Bulk Dyad-seq

For all bulk Dyad-seq experiments, 100 ng of purified genomic DNA was resuspended in 20 µL of glucosylation mix (1x CutSmart buffer (NEB, B7204S), 2.5x UDP-glucose, 10 U T4-BGT (NEB, M0357L)) and the samples were incubated at 37°C for 16 hours. Afterwards, 10 µL of protease mix (100 µg protease (Qiagen, 19155), 1x CutSmart buffer) was added to each sample, and the samples were heated to 50°C for 5 hours, 75°C for 20 minutes, and 80°C for 5 minutes. After this, processing of the samples differed based on the version of Dyad-seq used.

For M-M-Dyad-seq and M-H-Dyad-seq, 10 µL of MspJI digestion mix (2 U MspJI, 1x enzyme activator solution, 1x CutSmart buffer) was added to each sample and the samples were heated to 37°C for 5 hours, and 65°C for 20 minutes. Next, 1 µL of barcoded 1 µM double-stranded adapter was added. Then 9 µL of ligation mix (1.11x T4 ligase reaction buffer, 4.44 mM ATP (NEB, P0756L), 2000 U T4 DNA ligase (NEB, M0202M)) was added to each sample, and the samples were incubated at 16°C for 16 hours.

For H-M-Dyad-seq and H-H-Dyad-seq, 10 µL of AbaSI digestion mix (10 U AbaSI (NEB, R0665S), 1x CutSmart buffer) was added to each sample and the samples were incubated at 25°C for 2 hours, and then heated to 65°C for 20 minutes. Next, 1 µL of barcoded 1 µM double-stranded adapter was added. Then 9 µL of ligation mix (1.11x T4 ligase reaction buffer, 4.44 mM ATP (NEB, P0756L), 2000 U T4 DNA ligase (NEB, M0202M)) was added to each sample, and the samples were incubated at 16°C for 16 hours.

After ligation, up to three barcoded libraries of the same type were pooled and all Dyad-seq versions were subjected to a 1x AMPure XP bead cleanup (Beckman Coulter, A63881), and eluted in 40 µL of water.

M-M-Dyad-seq and H-M-Dyad-seq samples were then concentrated to a volume of 28 µL and subjected to nucleobase conversion using the NEBNext enzymatic methyl-seq conversion module (NEB, E7125S) according to the manufacturer’s recommendations except for performing the final elution step in 40 µL of water. For M-H-Dyad-seq and H-H-Dyad-seq samples, nucleobase conversion was performed using the NEBNext enzymatic methyl-seq conversion module similar to what has been described previously^11^. Briefly, samples were first concentrated to a volume of 17 µL. Then 4 µL of formamide (Sigma-Aldrich, F9037-100ML) was added and the samples were heated to 85°C for 10 minutes before being quenched on ice. APOBEC nucleobase conversion was performed as described by the manufacturer except for two minor changes. Samples were incubated at 37°C for 16 hours, and the final elution step was performed using 40 µL of water.

To all Dyad-seq versions, the nucleobase converted samples were subjected to one round of linear amplification. To do this, 9 µL of amplification mix was added (5.56x NEBuffer 2.1 (NEB, B7202S), 2.22 mM dNTPs (NEB, N0447L), and 2.22 uM linear amplification 9-mer: 5’-GCCTTGGCACCCGAGAATTCCANNNNNNNNN −3’) and the samples were heated to 95°C for 45 seconds before being quenched on ice. Once cold, 100 U of high concentration Klenow DNA polymerase (3’-5’ Exo-) (fisher scientific, 50-305-912) was added. Then samples were quickly vortexed, centrifuged and then incubated at 4°C for 5 minutes, followed by an increase of 1°C every 15 seconds at a ramp rate of 0.1°C per second till the samples reach 37°C which was then held for an additional 1.5 hours. Afterwards a 1.1x AMPure XP bead cleanup was performed, and the samplers were eluted in 40 µL of water before being concentrated down to 10 µL. The entire sample was then used in a linear PCR reaction by adding 15 µL of PCR mix (1.67x high-fidelity PCR mix (NEB, M0541L) and 0.67 µM Extended RPI primer: 5’-AATGATACGGCGACCACCGAGATCTACACGTTCAGAGTTCTACAGTCCGACGATCGGTGTAGTGGGTTTGG-3’) and performing PCR as follows: Initial denaturing at 98°C for 30 seconds, followed by 16 cycles of 98°C for 10 seconds, 59°C for 30 seconds, and 72°C for 30 seconds, and a final extension step at 72°C for 1 minute. Next, 5 µL of the linear PCR product was amplified further in a standard Illumina library PCR reaction, incorporating a uniquely indexed i7 primer. The remaining linear PCR product was stored at −20°C. To the final sequencing library, two 0.825x AMPure XP bead cleanups were performed with a final elution volume of 15 µL in water. The libraries were then quantified on an Agilent Bioanalyzer and Qubit fluorometer. Finally, libraries were subjected to paired-end 150 bp Illumina sequencing on a HiSeq platform.

### Bulk RNA-seq

Total RNA was extracted using TRIzol (Ambion, 15596018). 50 ng of total RNA was heated to 65°C for 5 minutes and returned to ice. Thereafter, it was combined with 9 uL of reverse transcription mix (20 U RNAseOUT (Invitrogen, 10777-019), 1.11x first strand buffer, 11.11 mM DTT, 0.56 mM dNTPs (NEB, N0447S), 100 U Superscript II (Invitrogen, 18064-071), and 25 ng of barcoded reverse transcription primer) and the sample was incubated at 42°C for 75 minutes, 4°C for 5 minutes, and 70°C for 10 minutes. Each replicate received a different barcoded reverse transcription primer. The reverse transcription primers used here have been described previously in Wangsanuwat *et al.*^45^. Afterwards, 50 µL of second strand synthesis mix (1.2x second strand buffer (Invitrogen, 10812-014), 0.24 mM dNTPs (NEB, N0447S), 4 U E.coli DNA Ligase (Invitrogen, 18052-019), 15 U E.coli DNA Polymerase I (Invitrogen, 18010-025), 0.8 U RNase H (Invitrogen, 18021-071)) was added to each sample and the samples were incubated at 16°C for 2 hours. The barcoded replicates were then pooled, and a 1x AMPure XP bead (Beckman Coulter, A63881) cleanup was performed, eluting in 30 µL of water, which was subsequently concentrated to 6.4 µL. The molecules were amplified with IVT and an Illumina sequencing library was prepared as described in CEL-seq2^36^. Libraries were sequenced on an Illumina HiSeq platform obtaining 150 bp reads from both ends.

### scDyad&T-seq

4 µL of Vapor-Lock (QIAGEN, 981611) was manually dispensed into each well of a 384-well plate using a 12-channel pipette. All downstream dispensing into 384-well plates were performed using the Nanodrop II liquid handling robot (BioNex Solutions). To each well, 100 nL of uniquely barcoded reverse transcription primers (7.5 ng/µL) containing 6 nucleotide UMI was added. The reverse transcription primers used here were previously described in Grun *et al.* with the exception that a UMI length of 6 was used^46^. Next, 100 nL of lysis buffer (0.175% IGEPAL CA-630, 1.75 mM dNTPs (NEB, N0447S), 1:1,250,000 ERCC RNA spike-in mix (Ambion, 4456740), and 0.19 U RNase inhibitor (Clontech, 2313A)) was added to each well. Single cells were sorted into individual wells of a 384-well plate using FACS and stored at - 80°C. To begin processing, plates were heated to 65°C for 3 minutes and returned to ice. Next, 150 nL of reverse transcription mix (0.7 U RNAseOUT (Invitrogen, 10777-019), 2.33x first strand buffer, 23.33 mM DTT, and 3.5 U Superscript II (Invitrogen, 18064-071)) was added to each well and the plates were incubated at 42°C for 75 minutes, 4°C for 5 minutes, and 70°C for 10 minutes. Thereafter, 1.5 µL of second strand synthesis mix (1.23x second strand buffer (Invitrogen, 10812-014), 0.25 mM dNTPs (NEB, N0447S), 0.14 U *E. coli* DNA Ligase (Invitrogen, 18052-019), 0.56 U *E. coli* DNA Polymerase I (Invitrogen, 18010-025), 0.03 U RNase H (Invitrogen, 18021-071)) was added to each well and the plates were incubated at 16°C for 2 hours. Following this step, 650 nL of protease mix (6 µg protease (Qiagen, 19155), 3.85x NEBuffer 4 (NEB, B7004S)) was added to each well, and the plates were heated to 50°C for 15 hours, 75°C for 20 minutes, and 80°C for 5 minutes. Next, 500 nL of glucosylation mix (1 U T4-BGT (NEB, M0357L), 6x UDP-glucose, 1x NEBuffer 4) was added to each well and the plates were incubated at 37°C for 16 hours. Thereafter, 500 nL of protease mix (2 µg protease, 1x NEBuffer 4) was added to each well, and the plates were incubated at 50°C for 3 hours, 75°C for 20 minutes, and 80°C for 5 minutes. Next, 500 nL of MspJI endonuclease mix (1x NEBuffer 4, 8x enzyme activator solution, 0.1 U MspJI (NEB, R0661L)) was added to each well and the plates were incubated at 37°C for 4.5 hours, and then heated to 65°C for 25 minutes. To each well, 280 nL of uniquely barcoded 250 nM unphosphorylated double-stranded Dyad-seq adapters were added. Next, 720 nL of ligation mix (1.39x T4 ligase reaction buffer, 5.56 mM ATP (NEB, P0756L), 140 U T4 DNA ligase (NEB, M0202M)) was added to each well, and the plates were incubated at 16°C for 16 hours. After ligation, uniquely barcoded reaction wells were pooled using a multichannel pipette, and the oil phase was discarded. The aqueous phase was incubated for 30 minutes with 1x AMPure XP beads (Beckman Coulter, A63881), and then subjected to standard bead cleanup with the DNA eluted in 30 µL of water. After vacuum concentrating the elute to 6.4 µL, *in vitro* transcription (IVT) was performed as previously described in the scAba-seq and scMspJI-seq protocols^4, 8^. RNA enrichment from the IVT product was performed as described previously with the following modifications^35^. The entire IVT product was used for enrichment, 4 µL of 1µM biotinylated polyA primer (5’-AAAAAAAAAAAAAAAAAAAAAAAA/3BioTEG/ −3’), and 8 µL of Dynabeads MyOne Streptavidin C1 beads (Invitrogen, 65001) were used and resuspended in 24 µL of 2x B&W solution after establishing RNase-free conditions. In addition, the supernatant was saved for additional processing.

The supernatant from the RNA enrichment process contains unamplified barcoded scDyad-seq DNA molecules. A 1x AMPure XP bead cleanup was performed by incubating the samples with beads for 30 minutes and eluting in 40 µL of water. Samples were then concentrated to 28 µL and nucleobase conversion was performed as described above for bulk M-M-Dyad-seq. Samples were then subjected to four rounds of linear amplification. The first round was the same as described for bulk Dyad-seq. In subsequent rounds, samples were first heated to 95°C for 45 seconds before being quenched on ice. Once cold, 5 µL of amplification mix was added (1x NEBuffer 2.1 (NEB, B7202S), 2 mM dNTPs (NEB, N0447L), 2 uM Linear amplification 9-mer, and 10 U of high concentration Klenow DNA polymerase (3’-5’ Exo-) (fisher scientific, 50-305-912)). Samples were then quickly vortexed, centrifuged and the same thermocycler conditions were used as in the first round of linear amplification. After 4 rounds of linear amplification, sequencing libraries were prepared the same way as described for bulk Dyad-seq. Finally, 150 bp paired-end Illumina sequencing was performed on a HiSeq platform. Sequencing metrics and associated metadata for each cell can be found in Supplementary File 1.

scDyad-seq is performed similar to scDyad&T-seq, except the initial reverse transcription and second strand synthesis steps are replaced with the equivalent volume of 1x NEBuffer 4. In addition, as the transcriptome is not captured, IVT is not performed and steps involving RNA enrichment and processing are omitted.

### Chip-seq data processing

The following published serum grown E14 ChIP datasets were used in this study (GEO accessions): GSM1000123 (H3K9ac), GSE74055 (H3K9me1 and H3K27ac), GSE23943 (H3K4me3, H3K9me3, H3K27me3, and H3K36me3), and GSE77420 (H3K9me2). For all these datasets, the processed data file was downloaded from GEO and further processed if needed. For GSE74055, the bigwigCompare tool on Galaxy (version 2.1.1.20160309.6) was used for 1 kb bins to identify enriched regions compared to the input data, with bins with a log2 enrichment score greater than 2 being considered enriched regions. For GSE23943, peak calling was performed using MACS2 on Galaxy, and the resulting narrow peaks file was used as enriched regions. For GSE77420, the enrichment score for H3K9me2 in serum grown conditions was compared to the input score within 2 kb bins. Regions were considered enriched if the H3K9me2 score was greater than the input score for both replicates. For all samples, contiguous enriched regions were combined into a single region. When applicable, enriched regions were converted from mm9 to mm10 using the UCSC genome browser LiftOver tool.

### Bulk RNA-seq analysis

Bulk RNA-seq data was processed as described previously with the following modification^35^. Reads were mapped to the RefSeq gene model based on the mouse genome release mm10, along with the set of 92 ERCC spike-in molecules (Ambion, 4456740).

DESeq2 was used for normalization and differential gene expression calling^47^. Gene expression differences between each condition were evaluated using adaptive shrinkage to adjust the log fold change observed^48^. For differential gene expression calling an adjusted p-value cutoff of 0.01 and a shrunken log fold change cutoff of 0.75 was used. For visualization and clustering, variance stabilizing transformation was performed and batch effects from different reverse transcription primer barcodes were removed using the removeBatchEffect function in the LIMMA package^49^.

### Dyad-seq analysis

Dyad-seq provides information on methylation or hydroxymethylation levels as well as information on 5mCpG or 5hmCpG maintenance levels. These two outputs of Dyad-seq were analyzed separately. To quantify 5mCpG maintenance levels, read 1 was trimmed to 86 nucleotides, and then exact duplicates were removed using Clumpify from BBTools. Next, reads containing the correct PCR amplification sequence and correct barcode were extracted. These reads were then trimmed using the default settings of TrimGalore. For mapping, Bismark was used in conjunction with Bowtie2 v2.3.5 to map to the mm10 build of the mouse genome^38^. For experiments using K562 cells, the hg19 build of the human genome was used. After mapping, Bismark was used to further deduplicate samples based on UMI, cell barcode and mapping location. For libraries that were prepared using MspJI, a custom Perl script was used to identify 5mC positions based on the cutting preference of MspJI, and the methylation status of the opposing cytosine in a CpG or CpHpG dyad context was inferred from the nucleobase conversion. For libraries that were prepared using AbaSI, a custom Perl script was used to identify 5hmC positions based on the cutting preference of AbaSI, and the methylation status of the opposing cytosine in a CpG dyad context was inferred from the nucleobase conversion. To quantify absolute methylation or hydroxymethylation levels, the cell barcode and UMI were transferred from read 1 to read 2. Read 1 was trimmed using TrimGalore in paired-end mode. The 5’ end of read 1 was clipped by 20 bases and the 3’ end of read 2 was hard clipped 34 bases after detection of the PCR amplification sequence CCACATCACCCAAACC to remove potential bias arising from enzymatic digestion and to avoid recounting unmethylated, methylated or hydroxymethylated cytosines detected at CpG dyads. The 5’ end of read 2 was clipped by 9 bases to minimize potential bias arising from the linear amplification random 9-mer primer. Similarly, the 3’ end of read 1 was also hard clipped 9 bases after the Illumina adapter was detected. Each read was mapped separately to mm10 using Bismark, and both the resulting sam files were deduplicated further using UMI, cell barcode and mapping location. The bismark_methylation_extractor tool was then used to extract the methylation status of detected cytosines. Next, a custom Perl code was used to demultiplex detected cytosines to the respective single cells based on the associated cell barcode. Thereafter, for cytosines detected in a CpG context, information from read 1 and read 2 were merged. Then, using UMIs, duplicate cytosine coverage resulting from overlapping paired-end reads or generated during the random priming step were deduplicated. Custom codes for analyzing Dyad-seq data and the accompanying documentation is provided with this work (Supplementary Software or github.com/alexchialastri/scDyad-T-seq). Cells for which less than 25,000 CpG sites were covered were discarded from downstream DNA methylation analysis. To cluster cells based on the methylome, hierarchical clustering was used and the optimal number of clusters was assigned using silhouette scores. Supplementary File 1 provides statistics of the methylome and transcriptome quantified in individual cells.

### scDyad&T-seq gene expression analysis

Read 2 was trimmed using the default settings of TrimGalore. After trimming, STARsolo (STAR aligner version 2.7.8a) was used to map the reads to mm10 using the gene annotation file from Ensembl. The reads were again mapped to mm10 using the transposable elements annotation file described in TEtranscripts^50^. Transcripts with the same UMI were deduplicated and genes or transposable elements that were not detected in at least one cell were removed from any downstream analysis. The combined counts from genes and transposable elements for each cell was considered the expression profile of that cell and was used in downstream analysis.

The standard analysis pipeline in Seurat (version 3.1.5) was used for single-cell RNA expression normalization and analysis^51^. Cells containing more than 500 genes and more than 2,000 unique transcripts were used for downstream analysis. The default NormalizeData function was used to log normalize the data. The top 1,000 most variable genes were used for making principal components and the elbow method was used to determine the optimal number of principle components for clustering. UMAP-based clustering was performed by running the following functions: FindNeighbors, FindClusters, and RunUMAP. To identify DEGs, the FindAllMarkers or FindMarkers function was used. The Wilcoxon rank sum test was used to classify a gene as differentially expressed, requiring a natural log fold change of at least 0.1 and an adjusted p-value of less than 0.05.

## Supporting information

Supplementary File 1: Sequencing metrics and metadata

Supplementary Software

## Acknowledgements

We thank members of the Dey lab for helpful discussions. We would like to thank Dr. Amander Clark for the E14 cell line. We would like to thank Jennifer Smith at the Biological Nanostructures Laboratory in the California NanoSystems Institute (CNSI), supported by UCSB and UC Office of the President, for help with Illumina sequencing. Computational work was supported by the Center for Scientific Computing at CNSI and Materials Research Laboratory (MRL) at UCSB: an NSF MRSEC (DMR-1720256) and NSF CNS-1725797. This work was supported by the NIH grants R01HD099517 and R01HG011013 to S.S.D.

## Author contributions

Conceptualization, A.C. and S.S.D.; Methodology, A.C. and S.S.D.; Investigation, A.C., S.S., E.E.S., and S.L.; Formal Analysis, A.C.; Writing – Original Draft, A.C.; Writing – Review & Editing, A.C. and S.S.D.; Funding Acquisition, S.S.D.; Resources, S.S.D.; Supervision, S.S.D.

## Data availability

Sequencing data have been deposited in the Gene Expression Omnibus (GEO) database accession code GEO: GSE197501.

## Code availability

Custom codes for analyzing Dyad-seq data and the accompanying documentation is provided with this work (Supplementary Software or github.com/alexchialastri/scDyad-T-seq).

