## Supplementary Information for "Combinatorial quantification of 5mC and 5hmC at individual CpG dyads and the transcriptome in single cells reveals modulators of DNA methylation maintenance fidelity"

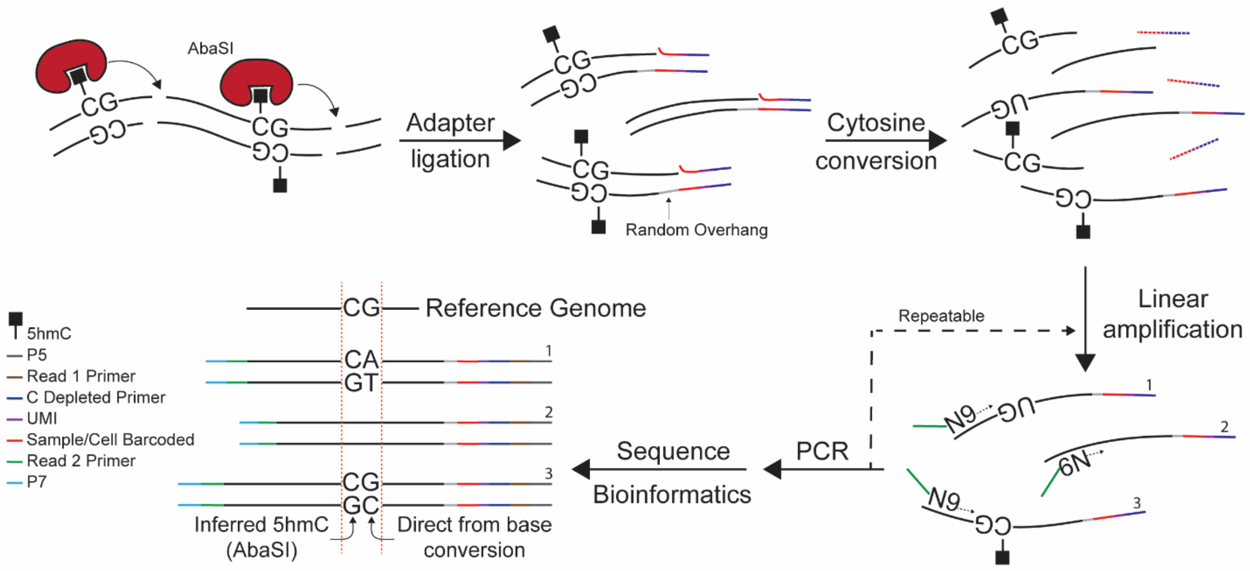


**Extended Data Figure 1 | Schematic of H-H-Dyad-seq.**

In H-H-Dyad-seq, the schematic shows that hydroxymethylated cytosines at CpG dyads are detected using AbaSI digestion followed by the appropriate nucleobase conversion to interrogate the hydroxymethylation status of the cytosine on the opposing strand of the dyad.

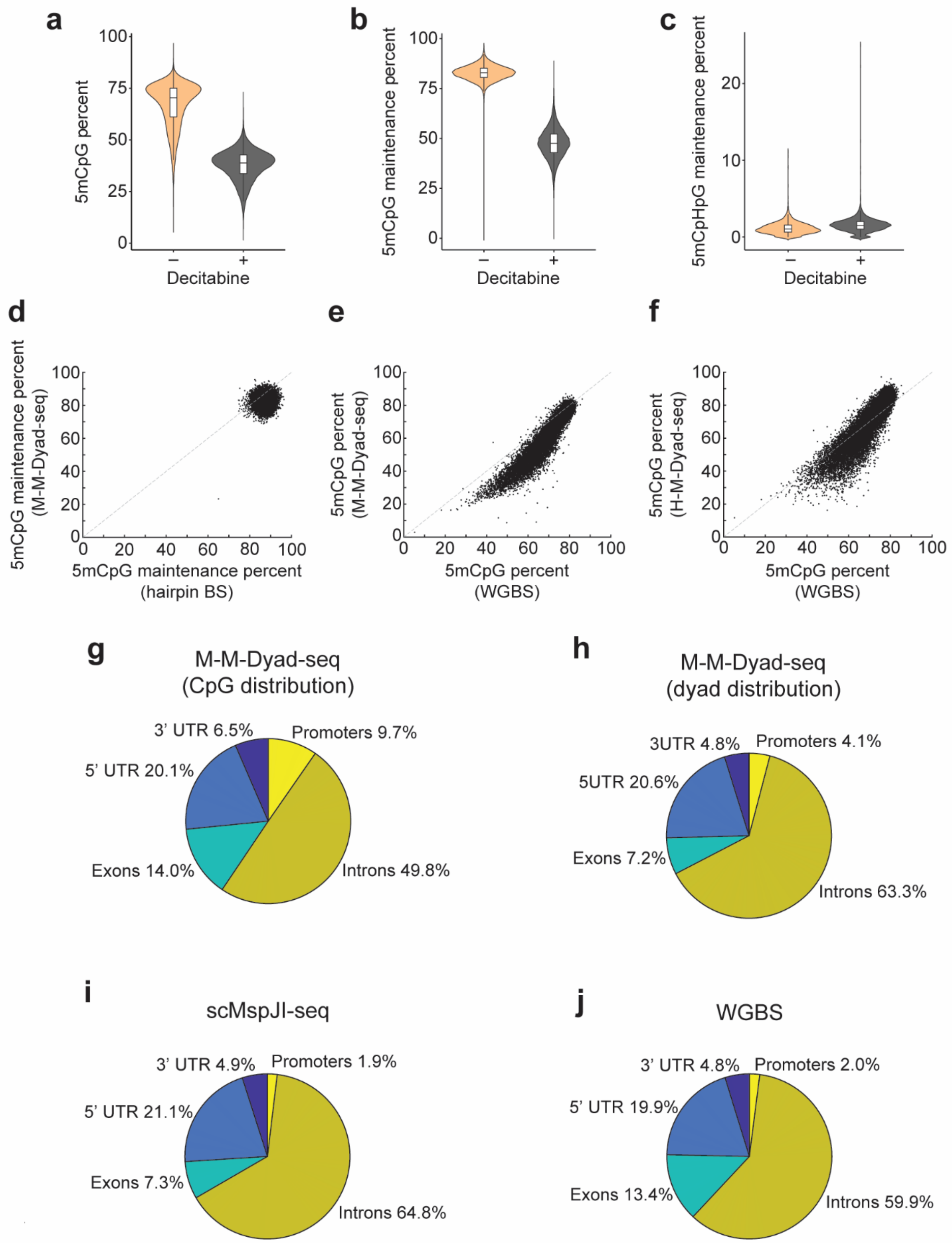


**Extended Data Figure 2 | M-M-Dyad-seq can accurately quantify DNA methylation levels and DNMT1-mediated maintenance methylation activity on a genome-wide scale.**

(**a**) M-M-Dyad-seq can quantify genome-wide methylation levels in mESCs grown under SL conditions in the absence (–) or presence (+) of the DNMT1 inhibitor Decitabine for 24 hours. (**b**)

M-M-Dyad-seq can also estimate genome-wide 5mCpG maintenance levels, quantified as the percentage of CpG dyads that are symmetrically methylated. Panels (a) and (b) show dramatic reduction in genome-wide methylation levels and 5mCpG maintenance levels in mESCs after treatment with Decitabine for 24 hours. (**c**) Panel shows 5mCpHpG maintenance methylation detected by M-M-Dyad-seq. (**d**) M-M-Dyad-seq shows similar 5mCpG maintenance levels as hairpin bisulfite sequencing in SL cultured mESCs[1]. Each point corresponds to a 100 kb bin the genome. (**e**) Genome-wide 5mCpG levels quantified by M-M-Dyad-seq correlates well with results obtained from bisulfite sequencing for SL cultured mESCs (Pearson $r$ = 0.94, Spearman $\rho$ = 0.94)[2]. Each point corresponds to a 100 kb bin the genome. (**f**) Genome-wide 5mCpG levels quantified by H-M-Dyad-seq correlates well with results obtained from bisulfite sequencing for SL cultured mESCs (Pearson $r$ = 0.90, Spearman $\rho$ = 0.90)[2]. Each point corresponds to a 100 kb bin the genome. (**g-j**) Pie chart shows that the distribution of 5mCpG sites detected over promoters, 5' UTRs, exons, introns and 3' UTRs is similar between M-M-Dyad-seq (panels g,h) and other techniques, such as scMspJI-seq (panel i) and whole-genome bisulfite sequencing (WGBS) (panel j)[3,4]. Data in these panels corresponds to SL cultured mESCs.

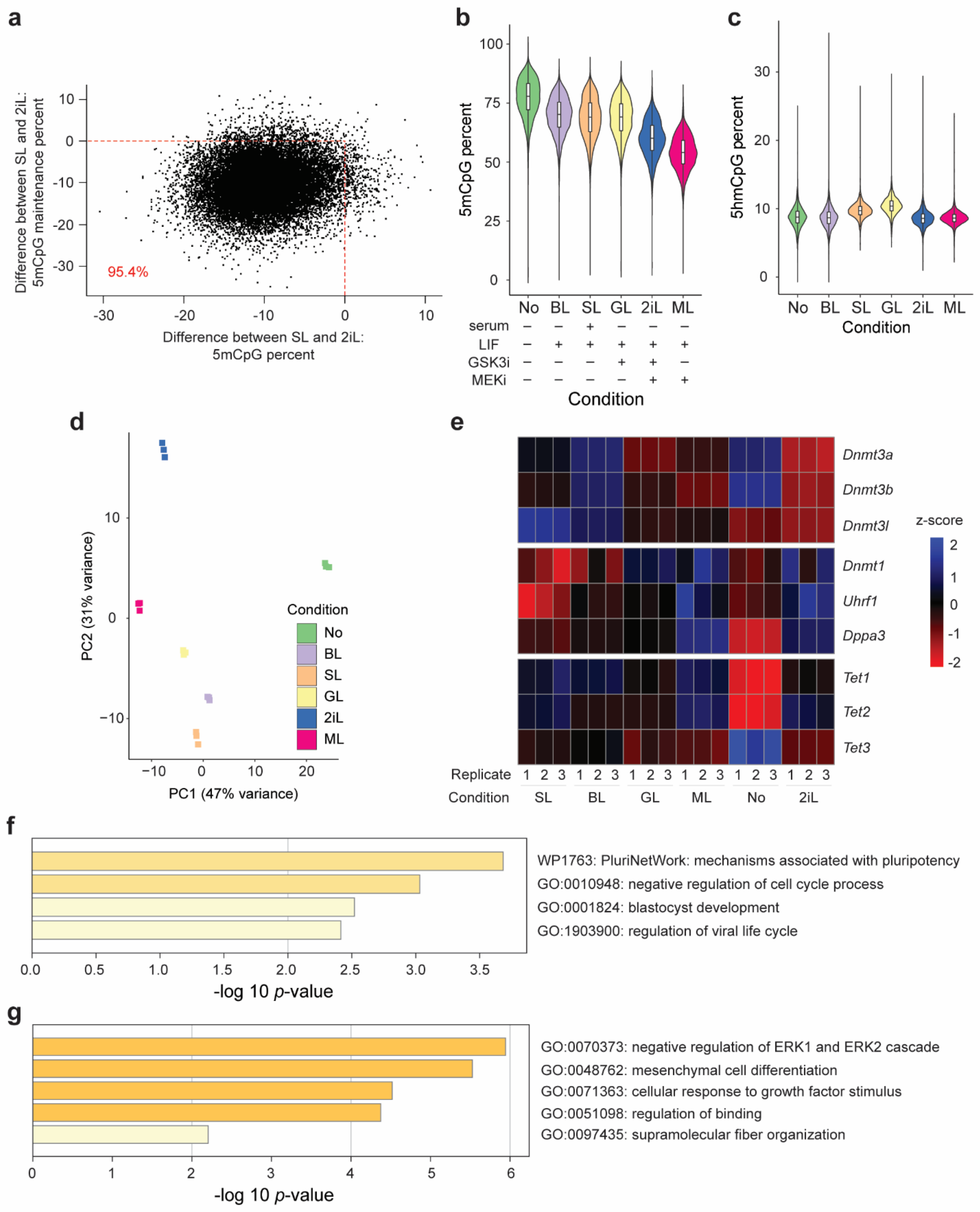


**Extended Data Figure 3 | Global DNA methylation and transcriptome reprogramming of mESCs transitioned from SL to different media conditions after 48 hours.**

(**a**) Loss of DNA methylation after culturing mESCs in 2iL conditions for 48 hours is associated with a reduction in 5mCpG maintenance levels. Each point represents genomic tilling of 100 kb. (**b**) Genome-wide 5mCpG levels quantified using H-M-Dyad-seq for mESCs grown under different conditions. (**c**) Genome-wide 5hmCpG levels quantified using H-H-Dyad-seq for mESCs grown under different conditions. (**d**) The first two principal components show distinct transcriptomes of mESCs grown under different conditions. Bulk RNA-seq was performed in triplicate. (**e**) Heatmap of expression level of genes related to *de novo* methylation, maintenance methylation, and demethylation pathways. (**f,g**) Gene pathway enrichment analysis for differentially expressed genes performed using Metascape[5]. Panel (f) shows gene sets associated with specific pathways that are highly expressed in 2iL and ML conditions, lowly expressed in No, and not differentially expressed across SL, BL, and GL conditions. Panel (g) shows gene sets associated with specific pathways that are highly expressed in the No condition, lowly expressed in 2iL and ML, and not differentially expressed across SL, BL, and GL.

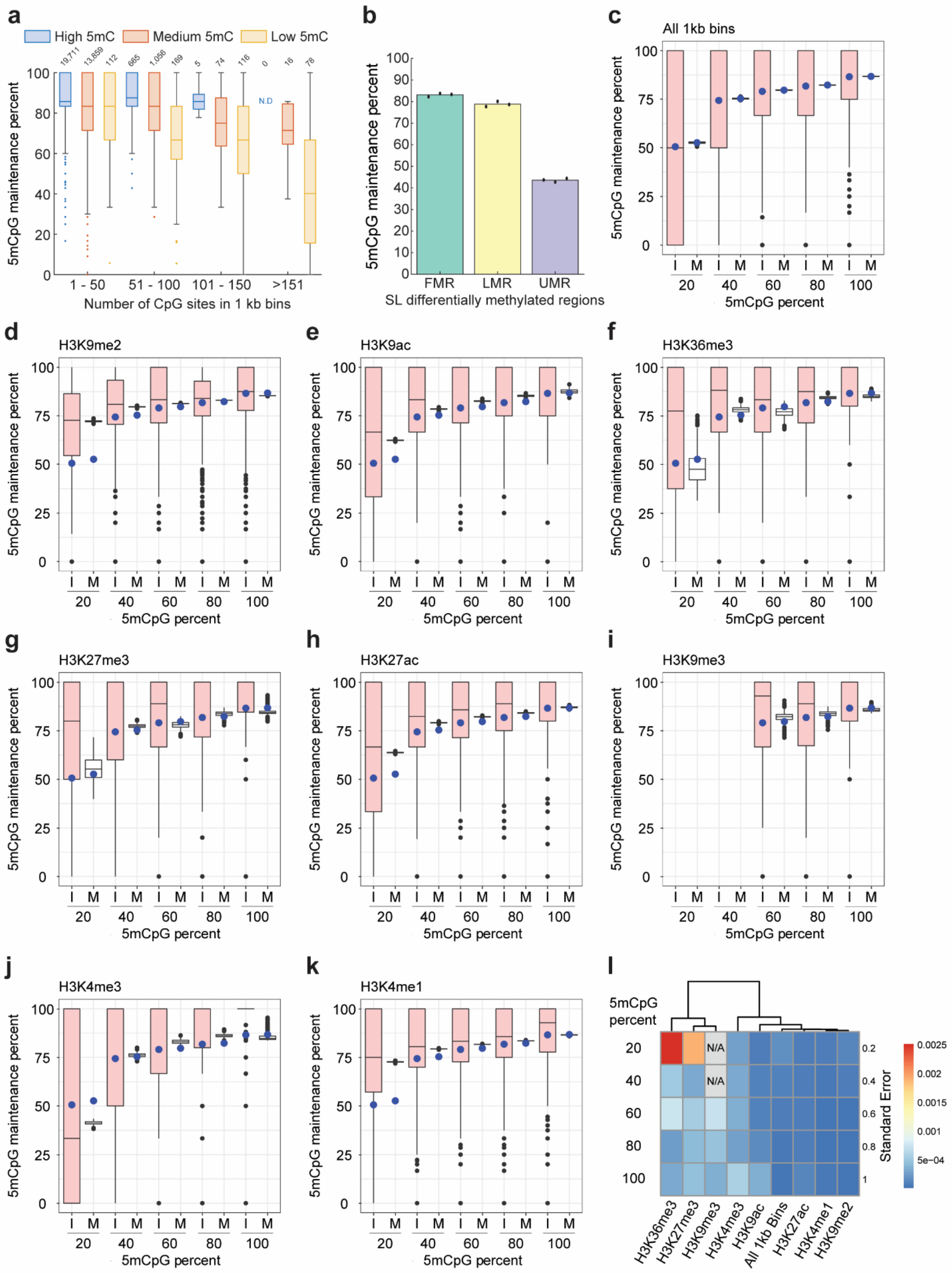


**Extended Data Figure 4 | Local density of the methylome and specific histone marks alter**

**5mCpG maintenance activity.**

(**a**) Box plot of 5mCpG maintenance levels in 1 kb genomic bins categorized based on the number of CpG sites in the bin and the absolute methylation levels. 'Low 5mC' indicates methylation levels lower than 20%, 'Medium 5mC' indicates methylation levels between 20% and 80%, and 'High 5mC' indicates methylation levels greater than 80%. N.D. stands for "Not detected". 1 kb regions in which at least 5 unique CpG dyads are detected were included in this panel. The number of bins in each category is denoted above each boxplot. Data in this panel corresponds to mESCs grown in SL condition and profiled using M-M-Dyad-seq. (**b**) 5mCpG maintenance levels at fully methylated regions (FMR), lowly methylated regions (LMR), and unmethylated regions (UMR) as stratified by Stadler *et al.*[6]. Data in this panel corresponds to mESCs grown in SL condition and profiled using M-M-Dyad-seq. (**c-k**) Box plots of 5mCpG maintenance levels as a function of absolute 5mCpG levels at individual loci enriched for a histone mark ('I') or a meta-region ('M') containing all enriched loci corresponding to a histone mark. Distributions for the meta-regions were obtained using bootstrapping, where resampling was performed 1,000 times per histone mark. Blue dots indicate average values found in genome-wide 1kb bins (same as data presented in panel (c)). (**l**) Standard error for the meta-regions in panels (c-k) and Figure 3e.

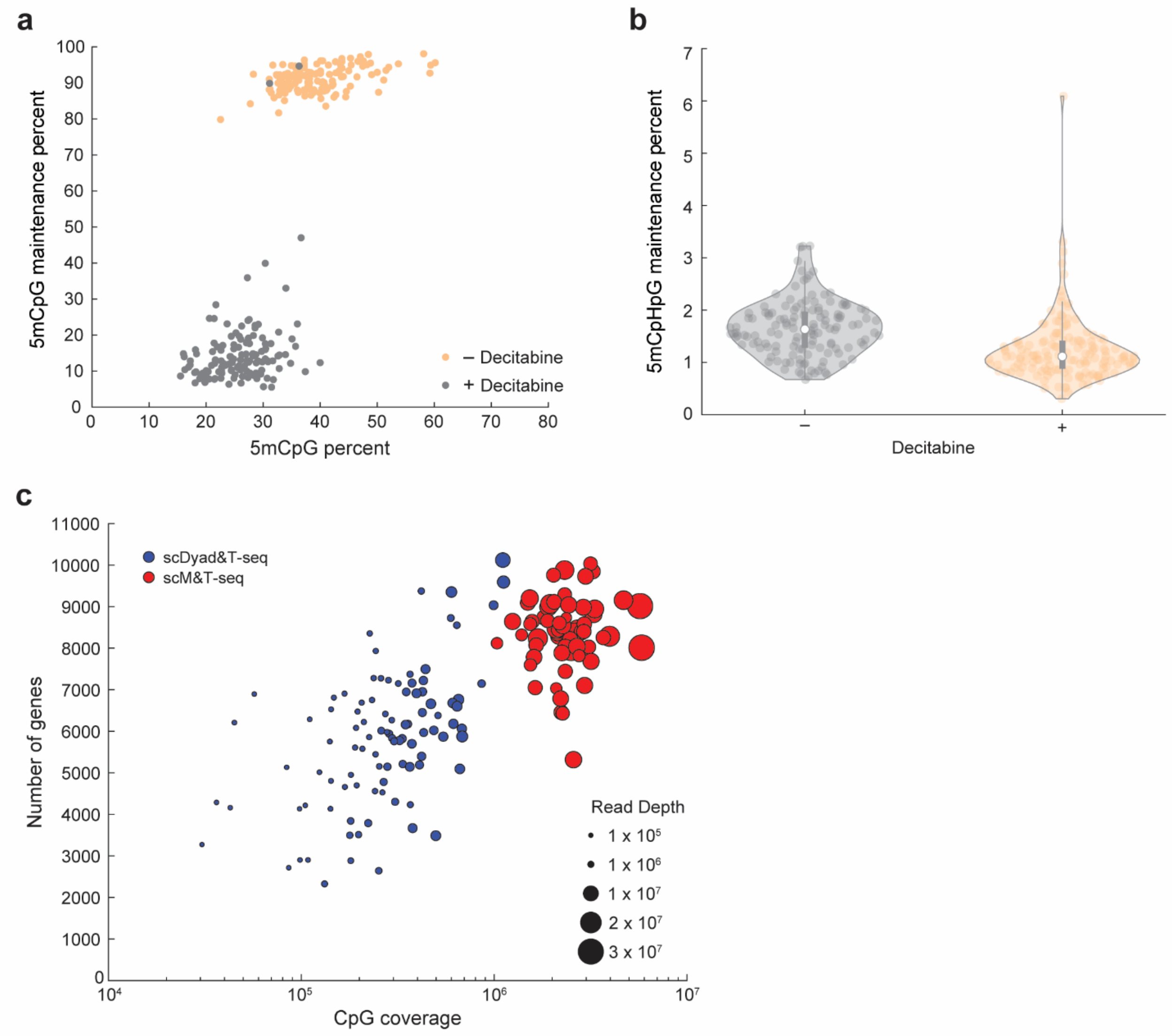


**Extended Data Figure 5 | scDyad&T-seq enables combined measurement of DNA methylation levels, 5mCpG maintenance levels and the transcriptome from the same cell.** (**a**) Genome-wide methylation and 5mCpG maintenance levels of single K562 cells treated with (+) or without (–) 0.6 μM Decitabine for 24 hours. (**b**) 5mCpHpG maintenance levels of single K562 cells treated with (+) or without (–) 0.6 μM Decitabine for 24 hours. (**c**) Coverage of CpG sites and the number of genes detected per cell in scDyad&T-seq (blue) and scM&T-seq (red)[7]. The diameter of each circle corresponds to the read depth at which a cell was sequenced. Data in this panel corresponds to SL cultured mESCs.

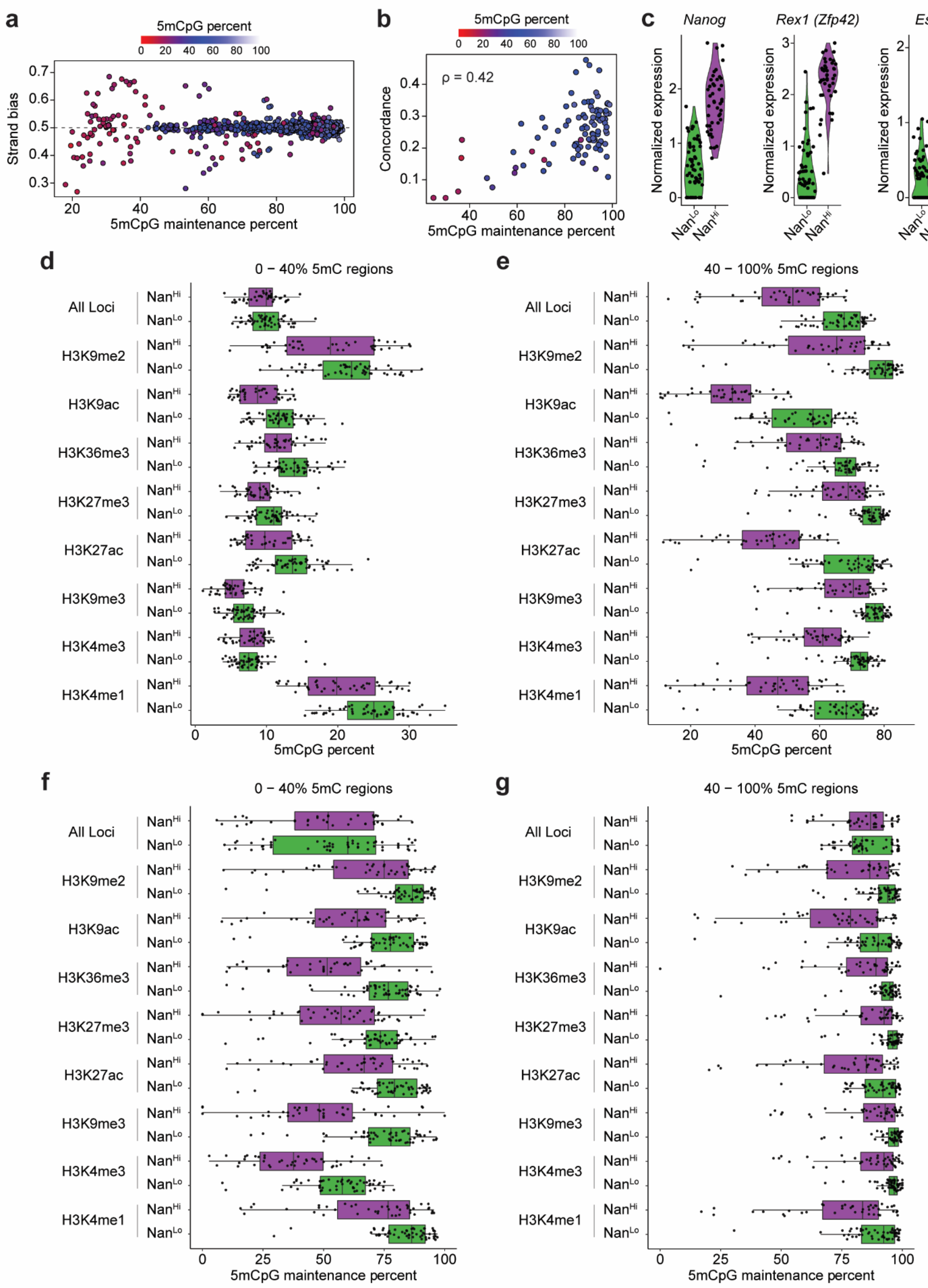


**Extended Data Figure 6 | Serum grown mESCs contain two distinct transcriptomic**

**subpopulations with Nanog high cells exhibiting decreased DNA methylation and 5mCpG maintenance levels across a broad range of histone modifications.**
(**a**) Comparison of chromosome-wide 5mCpG strand bias scores, estimated using techniques such as scMspJI-seq, to 5mCpG maintenance percent estimated using scDyad&T-seq. The color of the data points correspond to the absolute methylation levels estimated using scDyad&T-seq. (**b**) Comparison of genome-wide concordance of methylation calls to 5mCpG maintenance percent estimated using scDyad&T-seq for single cells[8]. Concordance is defined as the fraction of reads (with at least 5 CpG sites covered) where 90% or more of the sites are methylated. The color of the data points correspond to the absolute methylation levels estimated using scDyad&T-seq. (**c**) Expression level of pluripotency related genes NANOG, REX1, and ESRRB in the two transcriptional clusters (NANOG high ('$Nan^{Hi}$') and NANOG low ('$Nan^{Lo}$')) identified in Figure 5d using scDyad&T-seq for serum grown mESCs. (**d,e**) DNA methylation levels at regions marked by different histone modifications. (**f,g**) 5mCpG maintenance at regions marked by different histone modifications. From bulk measurements (see Fig. 3e), regions were previously categorized as less than (panels (d,f)) or greater than (panels (e,g)) 40% methylated.

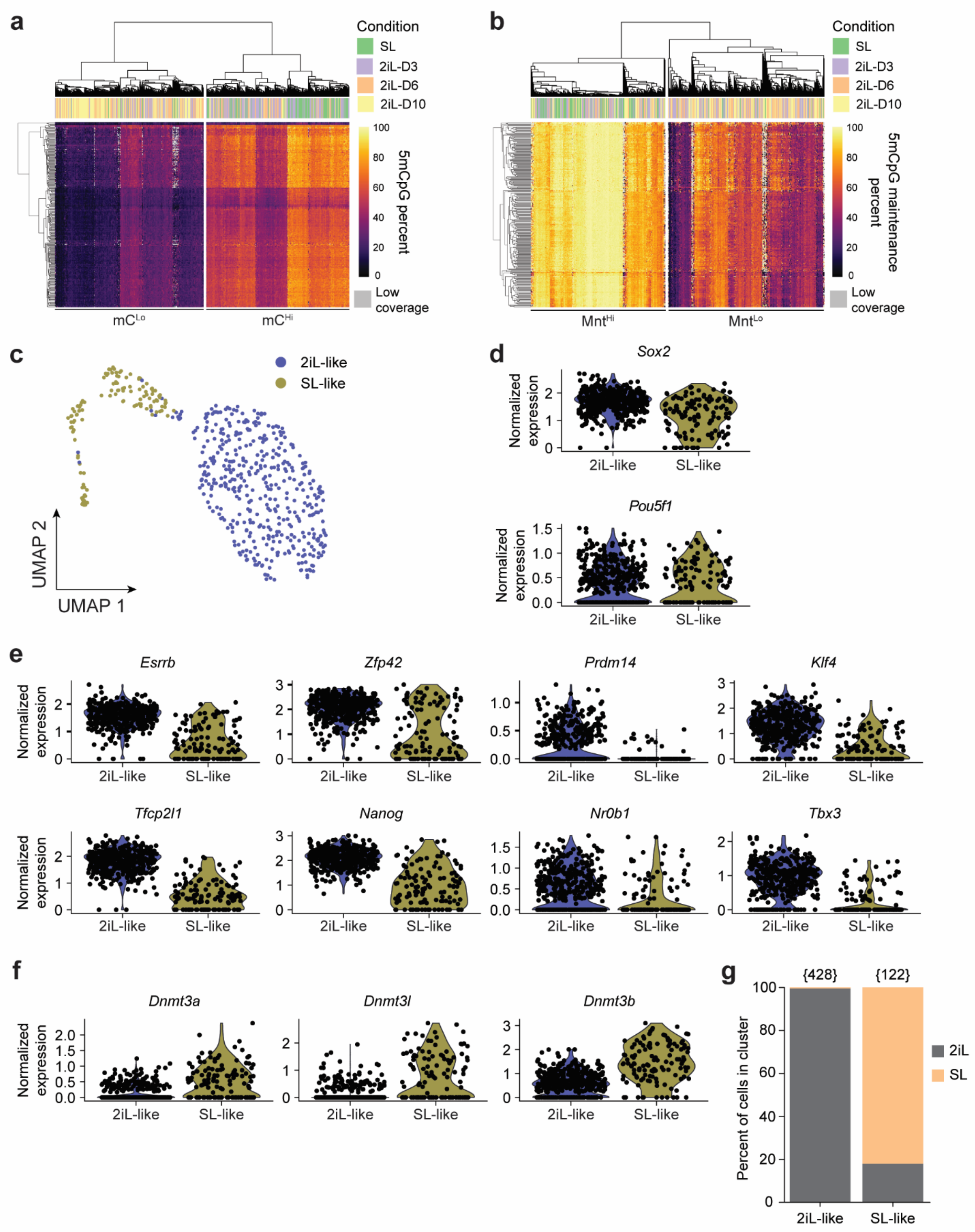


**Extended Data Figure 7 | Epigenetic and transcriptional reprogramming of mESCs transitioning from SL to 2iL conditions.**

(**a**) Hierarchical clustering based on genome-wide 5mCpG levels show that cells transitioning from SL to 2iL conditions can be classified into two major groups – a 5mCpG low ($mC^{Lo}$) or a 5mCpG high ($mC^{Hi}$) state. (**b**) Hierarchical clustering based on genome-wide 5mCpG maintenance levels show that cells transitioning from SL to 2iL conditions can be classified into two major groups – a lowly maintained ($Mnt^{Lo}$) or a highly maintained ($Mnt^{Hi}$) state. (**c**) UMAP visualization of cells transiting from SL to 2iL conditions, based on the single-cell transcriptomes obtained from scDyad&T-seq, show that cells can be classified into two broad transcriptional clusters. The cluster names, 2iL-like and SL-like were assigned based on expression of key marker genes in mESCs grown in 2iL or SL conditions, respectively. (**d**) Expression of key pluripotency genes that have previously been shown to be similar between SL and 2iL culture[9]. (**e**) Expression of genes known to be transcribed at higher levels in 2iL mESCs compared to those grown in SL conditions[9]. (**f**) Expression of genes known to be transcribed at higher levels in SL mESCs compared to those grown in 2iL culture[9]. (**g**) Bar plot shows the percentage of 2iL and SL grown mESCs that are assigned to the 2iL-like or SL-like transcriptional clusters. The number in the parenthesis indicates the total number of cells in that cluster.

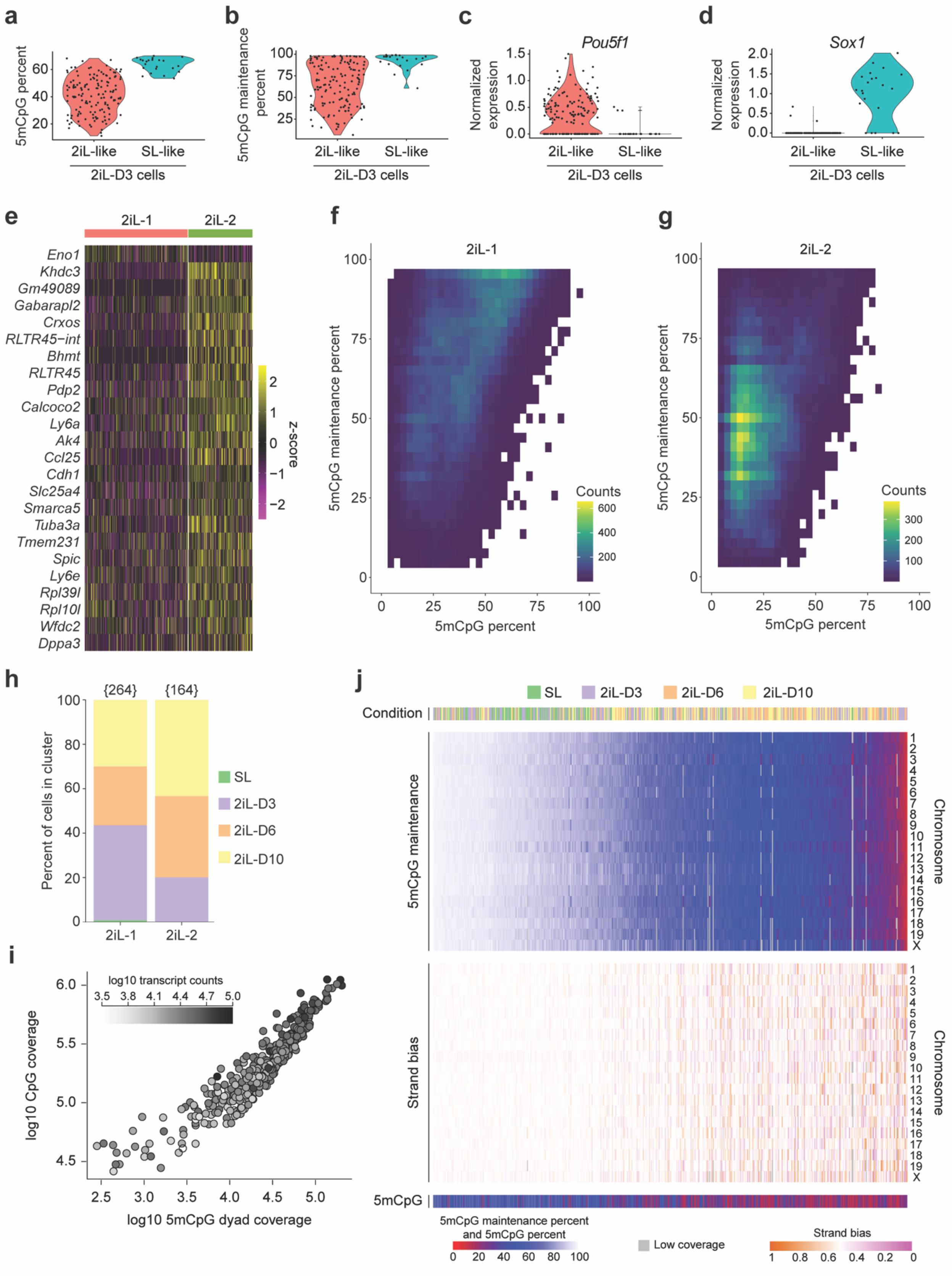


**Extended Data Figure 8 | scDyad&T-seq directly relates transcriptional cell identity to**

**demethylation dynamics in single cells transitioning from SL to 2iL.**

(**a,b**) Genome-wide DNA methylation (panel a) and 5mCpG maintenance (panel b) levels of 2iL-D3 cells in the two broad transcriptional groups – 2iL-like and SL-like – described in Extended Data Figure 7c. (**c,d**) Expression levels of the pluripotency marker POU5F1 (also known as OCT4) (panel c) and early neuroectoderm lineage marker SOX1 (panel d) in 2iL-D3 cells. (**e**) Heatmap of differentially expressed genes between the 2iL-1 and 2iL-2 population. (**f,g**) Absolute DNA methylation levels *vs.* the corresponding 5mCpG maintenance levels within 100 kb bins for cells in sub-population 2iL-1 (panel f) or 2iL-2 (panel g). (**h**) Bar plot shows how cells cultured in the 2iL condition for different number of days are distributed between the 2iL-1 and 2iL-2 sub-populations. The number in the parenthesis indicates the total number of cells in that sub-population. (**i**) Panel shows the coverage of CpG dinucleotides providing information on 5mCpG maintenance (dyad coverage), and the coverage of CpG sites providing information on the absolute levels of DNA methylation in single cells (CpG coverage). The color of the data points indicate the total number of unique transcripts detected in single cells grown in SL and 2iL conditions. (**j**) Heatmap of 5mCpG maintenance for individual chromosomes in single cells indicates increased sensitivity in quantifying DNMT1-mediated maintenance fidelity and demethylation compared to the strand bias score obtained from methods such as scMspJI-seq. The panel also shows the culture conditions and genome-wide 5mCpG methylation levels for the same cells.

**Supplementary Table 1 | Differentially expressed genes.** List of all genes that were found to be differentially expressed using scDyad&T-seq between the two subpopulations – $Nan^{Hi}$ and $Nan^{Lo}$ – in SL grown mESCs.

| Gene | Average lnFC | Percent of $NAN^{Lo}$ cells expressing gene | Percent of $NAN^{Hi}$ cells expressing gene | Adjusted p value | Subpopulation where gene is highly expressed |
|---|---|---|---|---|---|
| **Tubb6** | 1.01 | 98.30% | 79.50% | 1.63E-07 | $NAN^{Lo}$ |
| **Actg1** | 0.57 | 100.00% | 100.00% | 2.47E-06 | $NAN^{Lo}$ |
| **Car2** | 1.29 | 96.60% | 77.30% | 5.13E-06 | $NAN^{Lo}$ |
| **Cldn7** | 0.56 | 75.90% | 15.90% | 1.04E-05 | $NAN^{Lo}$ |
| **Zyx** | 0.76 | 86.20% | 31.80% | 1.10E-05 | $NAN^{Lo}$ |
| **Sema6a** | 0.52 | 65.50% | 2.30% | 1.11E-05 | $NAN^{Lo}$ |
| **Rap1gds1** | 0.42 | 81.00% | 20.50% | 1.20E-05 | $NAN^{Lo}$ |
| **H3f3b** | 0.45 | 100.00% | 100.00% | 1.75E-05 | $NAN^{Lo}$ |
| **Tuba1a** | 0.79 | 100.00% | 95.50% | 3.49E-05 | $NAN^{Lo}$ |
| **Pou3f1** | 0.76 | 74.10% | 13.60% | 4.47E-05 | $NAN^{Lo}$ |
| **Adgrl2** | 0.69 | 77.60% | 22.70% | 8.94E-05 | $NAN^{Lo}$ |
| **Arid2** | 0.6 | 89.70% | 54.50% | 9.09E-05 | $NAN^{Lo}$ |
| **Gnas** | 0.8 | 100.00% | 97.70% | 1.09E-04 | $NAN^{Lo}$ |
| **Rras2** | 0.72 | 86.20% | 40.90% | 1.69E-04 | $NAN^{Lo}$ |
| **Tpm4** | 0.88 | 91.40% | 65.90% | 1.96E-04 | $NAN^{Lo}$ |
| **Irs2** | 0.78 | 79.30% | 25.00% | 3.20E-04 | $NAN^{Lo}$ |
| **Nes** | 0.65 | 69.00% | 13.60% | 8.11E-04 | $NAN^{Lo}$ |
| **Rbpms** | 0.47 | 100.00% | 93.20% | 8.28E-04 | $NAN^{Lo}$ |
| **Gbp2b** | 0.89 | 60.30% | 6.80% | 9.33E-04 | $NAN^{Lo}$ |
| **Ssbp3** | 0.7 | 93.10% | 70.50% | 9.35E-04 | $NAN^{Lo}$ |
| **Cldn6** | 0.69 | 91.40% | 59.10% | 1.02E-03 | $NAN^{Lo}$ |
| **Arl4c** | 0.54 | 63.80% | 11.40% | 1.36E-03 | $NAN^{Lo}$ |
| **Perp** | 0.72 | 91.40% | 40.90% | 1.99E-03 | $NAN^{Lo}$ |
| **Peg10** | 1.06 | 98.30% | 86.40% | 2.02E-03 | $NAN^{Lo}$ |
| **Gbp2** | 0.8 | 58.60% | 6.80% | 2.08E-03 | $NAN^{Lo}$ |
| **Slc39a8** | 0.51 | 53.40% | 2.30% | 2.43E-03 | $NAN^{Lo}$ |
| **Abracl** | 0.59 | 93.10% | 54.50% | 2.52E-03 | $NAN^{Lo}$ |
| **Igf2bp3** | 0.58 | 91.40% | 70.50% | 2.58E-03 | $NAN^{Lo}$ |
| **Acsl3** | 0.62 | 84.50% | 47.70% | 2.85E-03 | $NAN^{Lo}$ |
| **Cald1** | 1.02 | 81.00% | 38.60% | 3.52E-03 | $NAN^{Lo}$ |
| **Crip2** | 0.56 | 91.40% | 65.90% | 3.94E-03 | $NAN^{Lo}$ |

| | | | | | |
|---|---|---|---|---|---|
| **Fndc3c1** | 0.31 | 55.20% | 4.50% | 4.11E-03 | $NAN^{Lo}$ |
| **Itgb1** | 0.65 | 96.60% | 84.10% | 4.62E-03 | $NAN^{Lo}$ |
| **Tagln** | 1.3 | 74.10% | 20.50% | 5.99E-03 | $NAN^{Lo}$ |
| **Cnn1** | 0.39 | 48.30% | 0.00% | 6.93E-03 | $NAN^{Lo}$ |
| **Dusp4** | 0.42 | 60.30% | 9.10% | 7.53E-03 | $NAN^{Lo}$ |
| **Ccng1** | 0.61 | 98.30% | 93.20% | 7.98E-03 | $NAN^{Lo}$ |
| **Tnfaip1** | 0.46 | 91.40% | 68.20% | 9.59E-03 | $NAN^{Lo}$ |
| **Igfbp3** | 0.88 | 67.20% | 18.20% | 9.69E-03 | $NAN^{Lo}$ |
| **Smo** | 0.31 | 67.20% | 13.60% | 9.89E-03 | $NAN^{Lo}$ |
| **Zcrb1** | 0.45 | 91.40% | 52.30% | 1.05E-02 | $NAN^{Lo}$ |
| **Myh10** | 0.46 | 89.70% | 59.10% | 1.17E-02 | $NAN^{Lo}$ |
| **Snrpb** | 0.37 | 100.00% | 95.50% | 1.26E-02 | $NAN^{Lo}$ |
| **Psmb8** | 0.44 | 50.00% | 2.30% | 1.59E-02 | $NAN^{Lo}$ |
| **Krt18** | 1.69 | 67.20% | 25.00% | 1.65E-02 | $NAN^{Lo}$ |
| **Pon2** | 0.32 | 60.30% | 11.40% | 1.75E-02 | $NAN^{Lo}$ |
| **Clu** | 0.58 | 70.70% | 25.00% | 2.07E-02 | $NAN^{Lo}$ |
| **Phgdh** | 0.36 | 98.30% | 79.50% | 2.09E-02 | $NAN^{Lo}$ |
| **Arhgdia** | 0.45 | 94.80% | 90.90% | 2.31E-02 | $NAN^{Lo}$ |
| **Cd24a** | 0.64 | 89.70% | 52.30% | 2.40E-02 | $NAN^{Lo}$ |
| **Bhlhe40** | 0.51 | 48.30% | 2.30% | 2.43E-02 | $NAN^{Lo}$ |
| **H2-K1** | 0.43 | 72.40% | 27.30% | 2.68E-02 | $NAN^{Lo}$ |
| **Rnf14** | 0.39 | 79.30% | 38.60% | 2.86E-02 | $NAN^{Lo}$ |
| **Plekha1** | 0.41 | 87.90% | 40.90% | 3.00E-02 | $NAN^{Lo}$ |
| **Prtg** | 0.65 | 72.40% | 27.30% | 3.11E-02 | $NAN^{Lo}$ |
| **Rap1b** | 0.41 | 98.30% | 93.20% | 3.35E-02 | $NAN^{Lo}$ |
| **Map2k4** | 0.45 | 91.40% | 56.80% | 3.38E-02 | $NAN^{Lo}$ |
| **Colec12** | 0.32 | 51.70% | 4.50% | 3.42E-02 | $NAN^{Lo}$ |
| **Ecpas** | 0.46 | 77.60% | 34.10% | 3.45E-02 | $NAN^{Lo}$ |
| **Vps36** | 0.4 | 94.80% | 70.50% | 3.67E-02 | $NAN^{Lo}$ |
| **Soat1** | 0.41 | 69.00% | 20.50% | 3.72E-02 | $NAN^{Lo}$ |
| **Vwa5a** | 0.37 | 63.80% | 15.90% | 4.17E-02 | $NAN^{Lo}$ |
| **Krt19** | 1.21 | 53.40% | 9.10% | 4.49E-02 | $NAN^{Lo}$ |
| **Irf1** | 0.56 | 72.40% | 29.50% | 4.74E-02 | $NAN^{Lo}$ |
| **Acadl** | 0.4 | 74.10% | 34.10% | 4.95E-02 | $NAN^{Lo}$ |
| **Zfp42** | 1.64 | 69.00% | 100.00% | 2.22E-11 | $NAN^{Hi}$ |
| **Nanog** | 1.21 | 77.60% | 100.00% | 2.78E-10 | $NAN^{Hi}$ |
| **Jam2** | 1.1 | 32.80% | 93.20% | 6.94E-10 | $NAN^{Hi}$ |
| **Esrrb** | 1.04 | 56.90% | 95.50% | 9.27E-10 | $NAN^{Hi}$ |
| **Morc1** | 0.81 | 29.30% | 88.60% | 2.31E-08 | $NAN^{Hi}$ |

| | | | | | |
|---|---|---|---|---|---|
| **L1Md-A** | 1.38 | 98.30% | 100.00% | 8.29E-08 | NAN$^{Hi}$ |
| **Stmn1** | 0.82 | 87.90% | 97.70% | 9.25E-08 | NAN$^{Hi}$ |
| **Mylpf** | 0.9 | 31.00% | 88.60% | 1.38E-07 | NAN$^{Hi}$ |
| **Sod2** | 0.71 | 77.60% | 97.70% | 3.37E-07 | NAN$^{Hi}$ |
| **Mtf2** | 0.67 | 96.60% | 100.00% | 4.04E-07 | NAN$^{Hi}$ |
| **Mybl2** | 0.77 | 79.30% | 97.70% | 6.16E-07 | NAN$^{Hi}$ |
| **Trap1a** | 0.72 | 55.20% | 90.90% | 1.45E-06 | NAN$^{Hi}$ |
| **AU018091** | 0.7 | 36.20% | 88.60% | 1.60E-06 | NAN$^{Hi}$ |
| **Fbxo15** | 0.98 | 48.30% | 86.40% | 1.89E-06 | NAN$^{Hi}$ |
| **Trim28** | 0.47 | 100.00% | 100.00% | 2.05E-06 | NAN$^{Hi}$ |
| **Lncenc1** | 0.73 | 32.80% | 84.10% | 3.39E-06 | NAN$^{Hi}$ |
| **Hmces** | 0.86 | 63.80% | 95.50% | 4.14E-06 | NAN$^{Hi}$ |
| **Zfp980** | 0.72 | 34.50% | 86.40% | 5.25E-06 | NAN$^{Hi}$ |
| **Epop** | 0.72 | 86.20% | 100.00% | 5.50E-06 | NAN$^{Hi}$ |
| **Hsd17b14** | 0.77 | 69.00% | 93.20% | 6.89E-06 | NAN$^{Hi}$ |
| **Chchd2** | 0.38 | 100.00% | 100.00% | 7.26E-06 | NAN$^{Hi}$ |
| **Spp1** | 0.87 | 31.00% | 86.40% | 9.74E-06 | NAN$^{Hi}$ |
| **RMER10A** | 0.71 | 81.00% | 95.50% | 1.13E-05 | NAN$^{Hi}$ |
| **Gsta4** | 0.75 | 87.90% | 97.70% | 1.19E-05 | NAN$^{Hi}$ |
| **Dppa5a** | 0.94 | 89.70% | 100.00% | 1.28E-05 | NAN$^{Hi}$ |
| **Msh6** | 0.56 | 98.30% | 100.00% | 1.68E-05 | NAN$^{Hi}$ |
| **Kat6b** | 0.68 | 50.00% | 88.60% | 1.68E-05 | NAN$^{Hi}$ |
| **Fgf4** | 0.66 | 37.90% | 86.40% | 1.74E-05 | NAN$^{Hi}$ |
| **Tdh** | 0.93 | 84.50% | 97.70% | 1.75E-05 | NAN$^{Hi}$ |
| **Gm46332** | 0.46 | 1.70% | 56.80% | 1.78E-05 | NAN$^{Hi}$ |
| **Eif2s2** | 0.66 | 100.00% | 100.00% | 2.08E-05 | NAN$^{Hi}$ |
| **Sgk1** | 0.75 | 53.40% | 93.20% | 2.67E-05 | NAN$^{Hi}$ |
| **Klf4** | 0.89 | 39.70% | 81.80% | 3.50E-05 | NAN$^{Hi}$ |
| **Epha4** | 0.67 | 29.30% | 79.50% | 4.17E-05 | NAN$^{Hi}$ |
| **Gm8935** | 0.69 | 46.60% | 90.90% | 4.41E-05 | NAN$^{Hi}$ |
| **Eprn** | 0.45 | 8.60% | 63.60% | 7.60E-05 | NAN$^{Hi}$ |
| **Cacybp** | 0.49 | 96.60% | 100.00% | 8.15E-05 | NAN$^{Hi}$ |
| **Nr0b1** | 0.72 | 19.00% | 70.50% | 8.23E-05 | NAN$^{Hi}$ |
| **RMER15** | 0.81 | 100.00% | 100.00% | 1.05E-04 | NAN$^{Hi}$ |
| **Rps4l** | 0.49 | 98.30% | 97.70% | 1.07E-04 | NAN$^{Hi}$ |
| **Calcoco2** | 0.89 | 20.70% | 70.50% | 1.85E-04 | NAN$^{Hi}$ |
| **Dnajc21** | 0.66 | 86.20% | 95.50% | 2.55E-04 | NAN$^{Hi}$ |
| **Lap3** | 0.59 | 94.80% | 100.00% | 2.93E-04 | NAN$^{Hi}$ |
| **1-Sep** | 0.35 | 3.40% | 54.50% | 3.34E-04 | NAN$^{Hi}$ |

| | | | | | |
|---|---|---|---|---|---|
| **Ifitm2** | 0.57 | 82.80% | 93.20% | 3.46E-04 | NAN$^{Hi}$ |
| **Enah** | 0.64 | 89.70% | 100.00% | 4.37E-04 | NAN$^{Hi}$ |
| **Rpl10l** | 0.66 | 27.60% | 72.70% | 5.96E-04 | NAN$^{Hi}$ |
| **Dnmt3l** | 0.73 | 51.70% | 95.50% | 6.59E-04 | NAN$^{Hi}$ |
| **Enox1** | 0.52 | 20.70% | 70.50% | 1.10E-03 | NAN$^{Hi}$ |
| **Slc28a1** | 0.52 | 15.50% | 63.60% | 1.16E-03 | NAN$^{Hi}$ |
| **Dhx16** | 0.59 | 75.90% | 90.90% | 1.17E-03 | NAN$^{Hi}$ |
| **Ooep** | 0.43 | 12.10% | 63.60% | 2.01E-03 | NAN$^{Hi}$ |
| **Hsp90aa1** | 0.33 | 100.00% | 100.00% | 2.03E-03 | NAN$^{Hi}$ |
| **Klf2** | 0.74 | 48.30% | 81.80% | 2.25E-03 | NAN$^{Hi}$ |
| **Mymx** | 0.28 | 0.00% | 43.20% | 2.45E-03 | NAN$^{Hi}$ |
| **Ulk1** | 0.59 | 50.00% | 84.10% | 2.81E-03 | NAN$^{Hi}$ |
| **Tigar** | 0.47 | 51.70% | 90.90% | 3.53E-03 | NAN$^{Hi}$ |
| **Zfp979** | 0.52 | 77.60% | 93.20% | 3.58E-03 | NAN$^{Hi}$ |
| **Zfp296** | 0.56 | 34.50% | 79.50% | 4.52E-03 | NAN$^{Hi}$ |
| **Ccnd3** | 0.52 | 91.40% | 100.00% | 4.83E-03 | NAN$^{Hi}$ |
| **Tfcp2l1** | 0.73 | 63.80% | 84.10% | 6.13E-03 | NAN$^{Hi}$ |
| **Rpl39l** | 0.71 | 8.60% | 54.50% | 6.19E-03 | NAN$^{Hi}$ |
| **Rps18** | 0.29 | 100.00% | 100.00% | 6.28E-03 | NAN$^{Hi}$ |
| **Sox2** | 0.53 | 87.90% | 97.70% | 6.60E-03 | NAN$^{Hi}$ |
| **Kdm3a** | 0.63 | 56.90% | 79.50% | 6.73E-03 | NAN$^{Hi}$ |
| **Chchd10** | 0.58 | 91.40% | 100.00% | 7.55E-03 | NAN$^{Hi}$ |
| **Zfp981** | 0.8 | 44.80% | 81.80% | 7.88E-03 | NAN$^{Hi}$ |
| **L1Md-T** | 0.52 | 100.00% | 100.00% | 8.44E-03 | NAN$^{Hi}$ |
| **Phf11d** | 0.34 | 12.10% | 59.10% | 1.08E-02 | NAN$^{Hi}$ |
| **Hsf2bp** | 0.41 | 19.00% | 65.90% | 1.16E-02 | NAN$^{Hi}$ |
| **Gm12346** | 0.5 | 82.80% | 95.50% | 1.37E-02 | NAN$^{Hi}$ |
| **Zfp534** | 0.5 | 50.00% | 81.80% | 1.43E-02 | NAN$^{Hi}$ |
| **Tcl1** | 0.46 | 32.80% | 70.50% | 2.08E-02 | NAN$^{Hi}$ |
| **B1-Mm** | 0.74 | 100.00% | 97.70% | 2.16E-02 | NAN$^{Hi}$ |
| **Ubxn2a** | 0.5 | 56.90% | 86.40% | 2.27E-02 | NAN$^{Hi}$ |
| **Carnmt1** | 0.45 | 86.20% | 100.00% | 2.39E-02 | NAN$^{Hi}$ |
| **Mras** | 0.49 | 17.20% | 61.40% | 2.39E-02 | NAN$^{Hi}$ |
| **Spink1** | 0.35 | 6.90% | 50.00% | 2.48E-02 | NAN$^{Hi}$ |
| **Slc25a12** | 0.54 | 48.30% | 77.30% | 2.50E-02 | NAN$^{Hi}$ |
| **Rbpj** | 0.51 | 84.50% | 95.50% | 2.63E-02 | NAN$^{Hi}$ |
| **Cdc5l** | 0.44 | 93.10% | 95.50% | 3.53E-02 | NAN$^{Hi}$ |
| **Ctnnal1** | 0.69 | 58.60% | 79.50% | 3.59E-02 | NAN$^{Hi}$ |
| **Hck** | 0.27 | 6.90% | 47.70% | 3.92E-02 | NAN$^{Hi}$ |

| | | | | | |
|---|---|---|---|---|---|
| **Trip12** | 0.38 | 98.30% | 95.50% | 4.06E-02 | $NAN^{Hi}$ |
| **Gm14820** | 0.2 | 0.00% | 36.40% | 4.24E-02 | $NAN^{Hi}$ |
| **Gdf3** | 0.48 | 63.80% | 95.50% | 4.48E-02 | $NAN^{Hi}$ |
| **Rps7** | 0.2 | 100.00% | 100.00% | 4.51E-02 | $NAN^{Hi}$ |
| **Zfp990** | 0.51 | 84.50% | 97.70% | 4.54E-02 | $NAN^{Hi}$ |
| **Smim40** | 0.34 | 24.10% | 63.60% | 4.96E-02 | $NAN^{Hi}$ |

**Supplementary Table 2 | Double-stranded adapter barcodes used in scDyad&T-seq.** Barcodes described 5' to 3' for the bottom adapter. Top adapter barcode sequences are the reverse compliment of those listed. Barcodes 1, 2 and 3 were used in bulk M-H-Dyad-seq.

| Barcode # | Cell barcode |
|---|---|
| 1 | AGAGATGGAA |
| 2 | TTGGATGGTA |
| 3 | TGTTTGTAGG |
| 4 | TGAAGAGAAG |
| 5 | AGTGTGAAGT |
| 6 | AAGTTGATGG |
| 7 | GAATGGTGAT |
| 8 | AGGTTGAGTT |
| 9 | GATTAGGTGA |
| 10 | GAAGATTGGT |
| 11 | AGAGGAAGAA |
| 12 | TGGTGTTATG |
| 13 | GAGAAGAAGA |
| 14 | GATATGGAAG |
| 15 | GTTAGGTAAG |
| 16 | GTTAGTAGAG |
| 17 | TGGTGTATGA |
| 18 | AGGAAAGTAG |
| 19 | GAGATAAAGG |
| 20 | TTGAAGGAGT |
| 21 | AGTGTGAGAA |
| 22 | GTAGATAGAG |
| 23 | AAGTGTTGAG |
| 24 | GTGAGTAGTT |
| 25 | AGAGGTTAGA |
| 26 | TGGTTGAAGA |
| 27 | ATGATGAAGG |
| 28 | AGTTGAGGAA |
| 29 | GAAAGTGATG |
| 30 | AAGGTTGGAA |
| 31 | GAATGGTATG |
| 32 | GAAGAGAAAG |
| 33 | AGGTTGAAAG |

| | |
|---|---|
| **34** | TAGGATGGAT |
| **35** | TAGGTGAAGT |
| **36** | ATGAGTGGAA |
| **37** | TAGATGTAGG |
| **38** | GATATGAGGT |
| **39** | GAAGAAAGTG |
| **40** | ATGAGAGTGA |
| **41** | AGGAGTATAG |
| **42** | GTGATAGATG |
| **43** | TGAGGTAGTA |
| **44** | GTGTGTAGAA |
| **45** | GAGAGTTGTA |
| **46** | GAGAAAGGTA |
| **47** | TTGATGGAAG |
| **48** | TGTATGGATG |
| **49** | AAGTAGGAAG |
| **50** | TGTGATGAAG |
| **51** | GATTGTGGTA |
| **52** | GATAGATAGG |
| **53** | GTATGGAAAG |
| **54** | GAGAAAGAAG |
| **55** | AGTGAAAGGA |
| **56** | TGATTGTGTG |
| **57** | TGAGATATGG |
| **58** | TAGTTTGAGG |
| **59** | AAGGTAGAAG |
| **60** | AGAGAGAAGA |
| **61** | GTTGGAAGAA |
| **62** | GAAGGATGTA |
| **63** | GTAAAGGAAG |
| **64** | ATGGAGAGTA |
| **65** | AGGAAGTTGT |
| **66** | TGATGGAGAT |
| **67** | TGAATGGTAG |
| **68** | GTTTGAAGGT |
| **69** | AGTTTGTGGT |
| **70** | AGATGTGAAG |
| **71** | AGATATGTGG |
| **72** | GAGAATAGTG |

| | |
|---|---|
| **73** | TTGGAAGAAG |
| **74** | TGGTTGTTAG |
| **75** | TGTTGAGATG |
| **76** | TAGGTTGTAG |
| **77** | GTGGTAAAGT |
| **78** | GAAGGAAAGA |
| **79** | GATAATGAGG |
| **80** | ATGTTGGTAG |
| **81** | GAGTGATAAG |
| **82** | AAGGTGAATG |
| **83** | TAGGAGTAAG |
| **84** | GTGAGATGAT |
| **85** | TGAGGTTAAG |
| **86** | TGAGAAAGGT |
| **87** | AGAGTTGATG |
| **88** | TTGGAAAGTG |
| **89** | GTTAGGAAGA |
| **90** | AGGATTGAAG |
| **91** | TAGAAGAGGT |
| **92** | GTGAAAGAGA |
| **93** | GTATGAGGAA |
| **94** | TGAAAGAGTG |
| **95** | GTTGTGAAAG |
| **96** | GAAAGAAGGT |

**Supplementary Table 3 | Prototype double-stranded adapter barcodes used in M-M-Dyad-seq.** Barcodes described 5’ to 3’ for the bottom adapter. Top adapter barcode sequences are the reverse compliment of those listed.

| Barcode # | Cell barcode |
|---|---|
| **1** | ATATGGAG |
| **2** | AGGGATTG |
| **86** | TGGTTGGA |

**Supplementary Table 4 | Double-stranded adapter barcodes used in H-M-Dyad-seq and H-H-Dyad-seq.** Barcodes described 5' to 3' for the bottom adapter. Top adapter barcode sequences are the reverse compliment of those listed.

| Barcode # | Cell barcode |
| --- | --- |
| **1** | GAGAATGTGT |
| **2** | AGGTAAGATG |
| **3** | TGTAAGTGAG |
