## Supplementary Software for "Combinatorial quantification of 5mC and 5hmC at individual CpG dyads and the transcriptome in single cells reveals modulators of DNA methylation maintenance fidelity": AbaSIFabaToCGtxt.pl

```
# Feature type	Chromosome/scaffold name	Start (bp)
$filename = $ARGV[0];
open($file, '<', $filename);

$outputfilenameZ      = substr($filename,0,-5)."_CGMaintained.txt";
open($outputZ, '>', $outputfilenameZ);

$outputfilenamez      = substr($filename,0,-5)."_CGNonMaintained.txt";
open($outputz, '>', $outputfilenamez);

while ($line = <$file>)
{
    chomp $line;
@x = split("\t",$line);

$x[0] = $x[0] + 1;
$x[1] = $x[1] + 1;
   # if ($x[3] == 1)
   # {
	#$x[2] = $x[2] + 1;
    #}
    #if ($x[3] == -1)
    #{
    #    $x[2] =$x[2] +3;
    #}
    
if ($x[8] eq Z) {print $outputZ ("$x[0] $x[1] $x[2] $x[4] $x[3]\n");}
if ($x[8] eq z) {print $outputz ("$x[0] $x[1] $x[2] $x[4] $x[3]\n");}

}
```
