## Supplementary Software for "Combinatorial quantification of 5mC and 5hmC at individual CpG dyads and the transcriptome in single cells reveals modulators of DNA methylation maintenance fidelity": MakeFastqShorter.pl

```
# Shorten a fastq file i.e. 150 read length to our normal 76
                                                                                                                
use warnings;

$Input = $ARGV[0];
$OutputName = $ARGV[1];
$LengthMake = $ARGV[2]; 

open($fastafileR1, $Input);
open($outputR1,'>',$OutputName);


while ($header = <$fastafileR1>)
{
    $sequence = <$fastafileR1>;
    $plus     = <$fastafileR1>;
    $qual     = <$fastafileR1>;
    chomp $header;
    chomp $sequence;
    chomp $plus;
    chomp $qual;
    $sequence2 = substr($sequence,0,$LengthMake);
    $qual2 = substr($qual,0,$LengthMake);
    
   # print $outputR1 ("$header$sequence2\n$plus$qual2\n");
    print $outputR1 ("$header\n$sequence2\n$plus\n$qual2\n");
	    
	
    
}
```
