## Supplementary Software for "Combinatorial quantification of 5mC and 5hmC at individual CpG dyads and the transcriptome in single cells reveals modulators of DNA methylation maintenance fidelity": process_sc5hmC_CpGDiad.pl

```
#!/usr/bin/perl

use strict "subs";
use strict "refs";
#use strict "vars";
 
print STDERR "
  processes sam file of single-cell 5hmC Dyad seq, adjusted from scAba-seq pipeline
 
  Usage
  process_scaba.pl genome_file.fa sam_file.sam aba_barcodes.csv
 
";

# read in genome file
my $genomefile = $ARGV[0];
open(MULTIFASTA, "$genomefile") || die " FATAL ERROR:\n Unable to load '$genomefile'.\n";
while (<MULTIFASTA>) {
	chomp($_);
	if ($_=~/^>(.*)/) {
		if ($seq) {
			$sequence{$header}=$seq;
		}
		$header = $1;
		$counter++;
		$seq    = '';
	} else {
		$seq.=$_;
	}
 
}
$sequence{$header}=$seq;

# read in cell-specific barcode file
my $file2 = $ARGV[2] or die "Need to get cell specific barcode file on the commend line\n";
my @csbc;
open(my $cscodes, '<', $file2) or die "Could not open '$file2' $!\n";
while (my $line = <$cscodes>) {
  chomp $line;
    my @fields = split ";" , $line;
    push @csbc,@fields[0];
}
my $BARCODELength = length($csbc[0]);

# read in sam file, filter reads, and count CG dinucleotides
my $file1 = $ARGV[1] or die "Need to get SAM file on the command line\n";
open(my $data, '<', $file1) or die "Could not open '$file1' $!\n";

my $UMILength = $ARGV[3] or die "Need UMI Length\n";

for ($i=0;$i<105;$i++)
 {
   $N_CG[$i] = 0;
 }
for ($k=0;$k<96;$k++)
 {
   $allreads[$k]=0;
   $rawreads[$k] = 0;
   $mappedreads[$k] = 0;
   $cleanreads[$k] = 0;
 }
 
 for ($k=0;$k<105;$k++){
					   $NucleotideA[$k]=0;
					   $NucleotideC[$k]=0;
					   $NucleotideG[$k]=0;
					   $NucleotideT[$k]=0;
					  
					 }
 
 
my $rABAoutputfile = substr($file1,0,-3)."raba";
my $fABAoutputfile = substr($file1,0,-3)."faba";
my @chrvec=qw(chr1 chr2 chr3 chr4 chr5 chr6 chr7 chr8 chr9 chr10 chr11 chr12 chr13 chr14 chr15 chr16 chr17 chr18 chr19 chr20 chr21 chr22 chrX chrY chrM);

open(my $abaout, '>', $rABAoutputfile) or die "Can't open file for writing: $!\n";

my $CGrABAoutputfile = substr($file1,0,-3)."CGraba";
my $CGfABAoutputfile = substr($file1,0,-3)."CGfaba";
my $CHGrABAoutputfile = substr($file1,0,-3)."CHGraba";
my $CHGfABAoutputfile = substr($file1,0,-3)."CHGfaba";

open(my $abaout, '>', $rABAoutputfile) or die "Can't open file for writing: $!\n";
open(my $CGabaout, '>', $CGrABAoutputfile) or die "Can't open file for writing: $!\n";
#open(my $CHGabaout, '>', $CHGrABAoutputfile) or die "Can't open file for writing: $!\n";


$N_plusmatch=0;
$N_minmatch=0;

$TotalUMINucleotides=0;
$CUMINucleotides=0;
$CGMaintained=0;
$CGNotMaintained=0;
#$CHGMaintained=0;
#$CHGNotMaintained=0;


while (my $line = <$data>) {
    chomp $line;
    my @fields = split "\t" , $line;
    my $seqflag = $fields[1];
    my $chr = $fields[2];
    my $cutcoord = $fields[3];
    my $cigar = $fields[5];

    my $uniqueflag = $fields[12];
    my $strand = 0;
    my $offset = 0;
    my $CGcount = 0;
 
 	my $fastqsequence = $fields[9];
	my $cellbarcode = substr($fields[0], index($fields[0], '-')+2+$UMILength,$BARCODELength);
	my $umi = substr($fields[0],index($fields[0], '-')+2,$UMILength);
	my $Methylationsequence = substr($fields[13],5);

	my $NCGDiad = 0;
	#my $NCHG = 0;
	
	#The methylation call string contains a dot ‘.’ for every position in the BS-read not involving a cytosine,
	# or contains one of the following letters for the three different cytosine methylation contexts (UPPER
	# CASE = METHYLATED, lower case = unmethylated):
	# z unmethylated C in CpG context
	# Z methylated C in CpG context
	# x unmethylated C in CHG context
	# X methylated C in CHG context
	# h unmethylated C in CHH context
	# H methylated C in CHH context
	# u unmethylated C in Unknown context (CN or CHN)
	# U methylated C in Unknown context (CN or CHN)
 
if ($cellbarcode ~~ @csbc ){
    my ($csbcindex) = grep {$csbc[$_] ~~ $cellbarcode } 0 .. 95;
	$allreads[$csbcindex]++;}

    if ($cellbarcode ~~ @csbc && $chr ~~ @chrvec){
	my ($csbcindex) = grep {$csbc[$_] ~~ $cellbarcode } 0 .. 95;
	$rawreads[$csbcindex]++;}
    if (($seqflag == 0 | $seqflag == 16) && ($cigar =~ /(^\d\d)M/) & (length($cigar)== 3) & $cellbarcode ~~ @csbc & $chr ~~ @chrvec){
       #$cigar eq "49M"
	   
	   my ($csbcindex) = grep {$csbc[$_] ~~ $cellbarcode } 0 .. 95;
       my ($chrindex) = grep {$chrvec[$_] ~~ $chr} 0 .. 24;       
       $mappedreads[$csbcindex]++;
       my $cutsite = uc(substr($sequence{$chr},$cutcoord-20,105));
	   
	   my $MatchLength = substr($cigar,0,2);
	   my $AdjustedLength = 70 - int($MatchLength); #70 was the read length the script was set up for (76 - 6 bc)
	   my $SecondAdjustLength = 49 - int($MatchLength); # 49 is the read length of 5mC CpG Dyad, This is to look up the methylation status of the Dyad
	   
	   
	   for ($k=0;$k<2;$k++){
		my $TempNucleotide = substr($umi,$k,1);
		$TotalUMINucleotides++;
	
		if ($TempNucleotide eq "C"){
		$CUMINucleotides++;
		}
	   }
	   
       my $char = 'CG';
       my $result = index($cutsite, $char, $offset); #will return the first instance of CG

       if ($result > -1) {$N_CG[$result]++; 
                                if ($result == 8  && $seqflag == 0){$CGpos = $cutcoord-12; $strand=1; $CGcount++}; #Indirect, 5hmC originally on + Strand
                                if ($result == 9  && $seqflag == 0){$CGpos = $cutcoord-11; $strand=1; $CGcount++}; #Indirect, 5hmC originally on + Strand (wobble)
								
								
                                if ($result == 29  && $seqflag == 0){$CGpos = $cutcoord+9; $strand=-1; $CGcount++;
								#Direct, 5hmC originally on - strand
								#$teststring =substr($cutsite,29,1);
								#$teststring =substr($fastqsequence,10,1); #Spot of the Opposite C
								$NCGDiad++;
								$NCGMaintained = substr ($Methylationsequence,10,1);
								}
								
                                if ($result == 30  && $seqflag == 0){$CGpos = $cutcoord+10; $strand=-1; $CGcount++;
								#Direct, 5hmC originally on - strand (wobble)
								$NCGDiad++;
								$NCGMaintained = substr ($Methylationsequence,11,1);
								} 
								
								
                                if ($result == 76-$AdjustedLength  && $seqflag == 16){$CGpos = $cutcoord+56-$AdjustedLength; $strand=1; $CGcount++;
								#Direct, 5hmC originally On + strand
								$NCGDiad++;
								$NCGMaintained = substr ($Methylationsequence,37-$SecondAdjustLength,1); #Mapping to - strand flips the read in sam file
								} 
								
                                if ($result == 77-$AdjustedLength  && $seqflag == 16){$CGpos = $cutcoord+57-$AdjustedLength; $strand=1; $CGcount++;
								#Direct, 5hmC originally On + strand (wobble)
								$NCGDiad++;
								$NCGMaintained = substr ($Methylationsequence,38-$SecondAdjustLength,1); #Mapping to - strand flips the read in sam file
								} 
								
								
                                if ($result == 97-$AdjustedLength  && $seqflag == 16){$CGpos = $cutcoord+77-$AdjustedLength; $strand=-1; $CGcount++}; #Indirect, 5hmC orignally on - strand
                                if ($result == 98-$AdjustedLength  && $seqflag == 16){$CGpos = $cutcoord+78-$AdjustedLength; $strand=-1; $CGcount++}; #Indirect, 5hmC orignally on - strand (wobble)
				$CGcand = uc(substr($sequence{$chr},$CGpos,2));
                                }

       while ($result != -1) { #this will keep marching down the list using the offset checking for the next CG site to assure there is only 1 CG in the right position and because there could randomly be a CG before the one we are looking for

          $offset = $result + 2;
          $result = index($cutsite, $char, $offset);
          if ($result > -1) {$N_CG[$result]++;

                                if ($result == 8  && $seqflag == 0){$CGpos = $cutcoord-12; $strand=1; $CGcount++};
                                if ($result == 9  && $seqflag == 0){$CGpos = $cutcoord-11; $strand=1; $CGcount++};
								
                                if ($result == 29  && $seqflag == 0){$CGpos = $cutcoord+9; $strand=-1; $CGcount++; $NCGDiad++; $NCGMaintained = substr ($Methylationsequence,10,1);}
                                if ($result == 30  && $seqflag == 0){$CGpos = $cutcoord+10; $strand=-1; $CGcount++; $NCGDiad++; $NCGMaintained = substr ($Methylationsequence,11,1);}
								
				if ($result == 76-$AdjustedLength  && $seqflag == 16){$CGpos = $cutcoord+56-$AdjustedLength; $strand=1; $CGcount++; $NCGDiad++; $NCGMaintained = substr ($Methylationsequence,37-$SecondAdjustLength,1); }
                                if ($result == 77-$AdjustedLength  && $seqflag == 16){$CGpos = $cutcoord+57-$AdjustedLength; $strand=1; $CGcount++; $NCGDiad++; $NCGMaintained = substr ($Methylationsequence,38-$SecondAdjustLength,1);}
								
                                if ($result == 97-$AdjustedLength  && $seqflag == 16){$CGpos = $cutcoord+77-$AdjustedLength; $strand=-1; $CGcount++};
                                if ($result == 98-$AdjustedLength  && $seqflag == 16){$CGpos = $cutcoord+78-$AdjustedLength; $strand=-1; $CGcount++};
                                $CGcand = uc(substr($sequence{$chr},$CGpos,2));
        			}}
	if ($CGcount==1){	
		print $abaout "$csbcindex\t$chrindex\t$CGpos\t$strand\t$umi\t$seqflag\t$CGcount\t$CGcand\t$NCGMaintained\n";
		$cleanreads[$csbcindex]++;}
	if ($CGcount==1 & $NCGDiad==1){
		if ($NCGMaintained eq 'Z'){$CGMaintained++;}
		elsif ($NCGMaintained eq 'z'){$CGNotMaintained++;}
		if ($NCGMaintained eq 'Z' | $NCGMaintained eq 'z') {
		print $CGabaout "$csbcindex\t$chrindex\t$CGpos\t$strand\t$umi\t$seqflag\t$CGcount\t$CGcand\t$NCGMaintained\n";
		}
		#print("$NCGMaintained \n");
		}
    }
} 
# write CG count to file
my $CGoutputfile = substr($file1,0,-3)."CG";
open(my $out, '>', $CGoutputfile) or die "Can't open file for writing: $!\n";
for ($i=0;$i<105;$i++)
{
    print $out "$N_CG[$i]\n";
}
# write QC count to file
my $QCoutputfile = substr($file1,0,-3)."QC";
open(my $qcout, '>', $QCoutputfile) or die "Can't open file for writing: $!\n";
my $QCoutputfile2 = substr($file1,0,-3)."QC2";
open(my $qcout2, '>', $QCoutputfile2) or die "Can't open file for writing: $!\n";

for ($k=0;$k<96;$k++)
 {
   print $qcout "$k \t $rawreads[$k] \t $mappedreads[$k] \t $cleanreads[$k] \n";
   print $qcout2 "$allreads[$k]\n";
 }
 
 #print basic maintenance Info
print "TotalUMINucleotides CUMINucleotides CGMaintained CGNotMaintained\n";
print "$TotalUMINucleotides $CUMINucleotides $CGMaintained $CGNotMaintained\n";

# keep only unique cuts and sort
system("sort -u -nk2 -nk3 -nk1 $rABAoutputfile > $fABAoutputfile");
system("sort -u $CGrABAoutputfile > $CGfABAoutputfile");
```
