## Supplementary Software for "Combinatorial quantification of 5mC and 5hmC at individual CpG dyads and the transcriptome in single cells reveals modulators of DNA methylation maintenance fidelity": process_scCpGDiadwithQC.pl

```
#!/usr/bin/perl

use strict "subs";
use strict "refs";
#use strict "vars";
 
print STDERR "
  processes sam file of single-cell MspJI-seq data
 
  Usage
  process_scaba.pl genome_file.fa sam_file.sam msjp1_barcodes.csv
 
";

# read in genome file
my $genomefile = $ARGV[0];
open(MULTIFASTA, "$genomefile") || die " FATAL ERROR:\n Unable to load '$genomefile'.\n";
while (<MULTIFASTA>) {
	chomp($_);
	if ($_=~/^>(.*)/) {
		if ($seq) {
			$sequence{$header}=$seq;
		}
		$header = $1;
		$counter++;
		$seq    = '';
	} else {
		$seq.=$_;
	}
 
}
$sequence{$header}=$seq;

# read in cell-specific barcode file
my $file2 = $ARGV[2] or die "Need to get cell specific barcode file on the commend line\n";
my @csbc;
open(my $cscodes, '<', $file2) or die "Could not open '$file2' $!\n";
while (my $line = <$cscodes>) {
  chomp $line;
    my @fields = split "\t" , $line;
    push @csbc,@fields[0];
}
my $BARCODELength = length($csbc[0]);

# read in sam file, filter reads, and count CG dinucleotides
my $file1 = $ARGV[1] or die "Need to get SAM file on the command line\n";
open(my $data, '<', $file1) or die "Could not open '$file1' $!\n";

my $UMILength = $ARGV[3] or die "Need UMI Length\n";

# for QC file
for ($k=0;$k<96;$k++)
 {
   $allreads[$k]=0;
   $rawreads[$k] = 0;
   $mappedreads[$k] = 0;
   $cleanreads[$k] = 0;
 }
 
					for ($k=0;$k<105;$k++)
					 {
					   $NucleotideA[$k]=0;
					   $NucleotideC[$k]=0;
					   $NucleotideG[$k]=0;
					   $NucleotideT[$k]=0;
					  
					 }

my $rABAoutputfile = substr($file1,0,-3)."raba";
my $fABAoutputfile = substr($file1,0,-3)."faba";
my $rABAoutputfileZymoSpikeIn = substr($file1,0,-3)."ZymoSpikeInraba";
my $fABAoutputfileZymoSpikeIn = substr($file1,0,-3)."ZymoSpikeInfaba";
my $rABAoutputfileLamndaPhageSpikeIn = substr($file1,0,-3)."LamndaPhageSpikeInraba";
my $fABAoutputfileLamndaPhageSpikeIn = substr($file1,0,-3)."LamndaPhageSpikeInfaba";
my @chrvec=qw(chr1 chr2 chr3 chr4 chr5 chr6 chr7 chr8 chr9 chr10 chr11 chr12 chr13 chr14 chr15 chr16 chr17 chr18 chr19 chr20 chr21 chr22 chrX chrY chrM ZymoDNAStandardSet PhageLambda);

my $CGrABAoutputfile = substr($file1,0,-3)."CGraba";
my $CGfABAoutputfile = substr($file1,0,-3)."CGfaba";
my $CHGrABAoutputfile = substr($file1,0,-3)."CHGraba";
my $CHGfABAoutputfile = substr($file1,0,-3)."CHGfaba";

open(my $abaout, '>', $rABAoutputfile) or die "Can't open file for writing: $!\n";
open(my $CGabaout, '>', $CGrABAoutputfile) or die "Can't open file for writing: $!\n";
open(my $CHGabaout, '>', $CHGrABAoutputfile) or die "Can't open file for writing: $!\n";
open(my $Zymoabaout, '>', $rABAoutputfileZymoSpikeIn) or die "Can't open file for writing: $!\n";
open(my $Lamndaabaout, '>', $rABAoutputfileLamndaPhageSpikeIn) or die "Can't open file for writing: $!\n";

$N_plusmatch=0;
$N_minmatch=0;

$TotalUMINucleotides=0;
$CUMINucleotides=0;
$CGMaintained=0;
$CGNotMaintained=0;
$CHGMaintained=0;
$CHGNotMaintained=0;
				
while (my $line = <$data>) {
    chomp $line;
    my @fields = split "\t" , $line;
    my $seqflag = $fields[1];
    my $chr = $fields[2];
    my $cutcoord = $fields[3];
    my $cigar = $fields[5];
   
    my $uniqueflag = $fields[12];
    my $strand = 0;
    my $offset = 0;
    my $CGcount = 0;
	
	my $fastqsequence = $fields[9];
	my $cellbarcode = substr($fields[0], index($fields[0], '-')+2+$UMILength,$BARCODELength);
	my $umi = substr($fields[0],index($fields[0], '-')+2,$UMILength);
	my $Methylationsequence = substr($fields[13],5);
    #print ("$cellbarcode $umi\n");
	my $NCG = 0;
	my $NCHG = 0;
	#The methylation call string contains a dot ‘.’ for every position in the BS-read not involving a cytosine,
	# or contains one of the following letters for the three different cytosine methylation contexts (UPPER
	# CASE = METHYLATED, lower case = unmethylated):
	# z unmethylated C in CpG context
	# Z methylated C in CpG context
	# x unmethylated C in CHG context
	# X methylated C in CHG context
	# h unmethylated C in CHH context
	# H methylated C in CHH context
	# u unmethylated C in Unknown context (CN or CHN)
	# U methylated C in Unknown context (CN or CHN)
    if ($cellbarcode ~~ @csbc ){

	$allreads[$csbcindex]++;}

    if ($cellbarcode ~~ @csbc & $chr ~~ @chrvec){
	my ($csbcindex) = grep {$csbc[$_] ~~ $cellbarcode } 0 .. 95;
	$rawreads[$csbcindex]++;}
    if (($seqflag == 0 | $seqflag == 16) & ($cigar =~ /(^\d\d)M/) & (length($cigar)== 3) & $cellbarcode ~~ @csbc & $chr ~~ @chrvec){
					
					#$cigar eq "49M"
	
	   
	   my ($csbcindex) = grep {$csbc[$_] ~~ $cellbarcode } 0 .. 95;
	  
       my ($chrindex) = grep {$chrvec[$_] ~~ $chr} 0 .. 26;       
       $mappedreads[$csbcindex]++;
       my $cutsite = uc(substr($sequence{$chr},$cutcoord-20,105));
       my $Nhits=0;
	   
	   my $MatchLength = substr($cigar,0,2);
       my $AdjustedLength = 65 - int($MatchLength); #65 was the read length the script was set up for (76 - 8 bc - 3 UMI)
	   my $SecondAdjustLength = 49 - int($MatchLength); # script developed for 49 bp in length then this adjusts for real mapping length 5mC CpG Dyad

	   my $SpikeInFlag=0; #These 3 lines create a flag for if a spike in was detected so that it can be put in it own file
	   if ($chrindex == 25){$SpikeInFlag=1;}
	   elsif($chrindex == 26){$SpikeInFlag=2;}
	   $NCGMaintained="N/A";$NCHGMaintained="N/A"; #Gives a standard value to these unless changed by the script below

	   for ($k=0;$k<4;$k++){
		my $TempNucleotide = substr($umi,$k,1);
		$TotalUMINucleotides++;
	
		if ($TempNucleotide eq "C"){
		$CUMINucleotides++;
		}
	   }
	   
	   
	   for ($k=0;$k<48;$k++){
					# if ($seqflag == 16 & substr($cutsite,51,1) eq 'C')
					 # {
					 #if (substr($cutsite,34,1) eq 'C'){
					 my $TempNucleotide = substr($fastqsequence,$k,1);
					 
					 if ($TempNucleotide eq "A") {$NucleotideA[$k]++;}
					 if ($TempNucleotide eq "C") {$NucleotideC[$k]++;}
					 if ($TempNucleotide eq "G") {$NucleotideG[$k]++;}
					 if ($TempNucleotide eq "T") {$NucleotideT[$k]++;}
					   
					  
					 #}
					 #}
					 }
	   
	   
	   
	   #Indirect, 5mC originally on + Strand
         if ($seqflag == 0 & substr($cutsite,6,1) eq 'C'){
	     $CGpos=$cutcoord-14; $CGcand = uc(substr($sequence{$chr},$CGpos,4));$strand=1; $Nhits++;
		 }
         
		 #Direct, 5mC originally On + strand
		 if ($seqflag == 16 & substr($cutsite,67-$AdjustedLength,1) eq 'C'){
		 $CGpos=$cutcoord+47-$AdjustedLength; $CGcand = uc(substr($sequence{$chr},$CGpos,4));$strand=1; $Nhits++;
		 			if (substr($cutsite,52-$SecondAdjustLength,1) eq 'G'){
						my $TempNucleotide = substr($fastqsequence,33-$SecondAdjustLength,1);
						#print "$TempNucleotide";
						my $TempMethStatus = substr ($Methylationsequence,33-$SecondAdjustLength,1);
						$CGpos2=$cutcoord+47-$AdjustedLength; $CGcand2 = uc(substr($sequence{$chr},$CGpos2,4));$strand2=1; $NCG++;
						if ($TempMethStatus eq 'Z'){
						$CGMaintained++;
						$NCGMaintained="Z";}
						elsif ($TempMethStatus eq 'z'){
						$CGNotMaintained++;
						$NCGMaintained="z";
						}
						else {$NCGMaintained=".";}
					}
					elsif (substr($cutsite,53-$SecondAdjustLength,1) eq 'G') {
					my $TempMethStatus = substr ($Methylationsequence,34-$SecondAdjustLength,1);
					$CGpos2=$cutcoord+47-$AdjustedLength; $CGcand2 = uc(substr($sequence{$chr},$CGpos2,4));$strand2=1; $NCHG++;
					if ($TempMethStatus eq 'X'){
					$CHGMaintained++;
					$NCHGMaintained="X";}
					elsif ($TempMethStatus eq 'x'){
					$CHGNotMaintained++;
					$NCHGMaintained="x";}
					else {$NCHGMaintained=".";}
					}
					
			}
		 
		 #Direct, 5mC originally on - strand
		 if ($seqflag == 0 & substr($cutsite,35,1) eq 'G'){
		$CGpos=$cutcoord+12; $CGcand = uc(substr($sequence{$chr},$CGpos,4)); $seqmatch = reverse $CGcand; $seqmatch =~ tr/ACGTacgt/TGCAtgca/; $CGcand = $seqmatch; $strand=-1; $Nhits++;
			if (substr($cutsite,34,1) eq 'C'){
				my $TempMethStatus = substr ($Methylationsequence,15,1);
				$CGpos2=$cutcoord+12; $CGcand2 = uc(substr($sequence{$chr},$CGpos,4)); $seqmatch2 = reverse $CGcand2; $seqmatch2 =~ tr/ACGTacgt/TGCAtgca/; $CGcand2 = $seqmatch2; $strand2=-1; $NCG++;

				if ($TempMethStatus eq 'Z'){
				$CGMaintained++;
				$NCGMaintained="Z";}
				elsif ($TempMethStatus eq 'z'){
				$CGNotMaintained++;
				$NCGMaintained="z";}
				else {$NCGMaintained=".";}
				}
				
			elsif (substr($cutsite,33,1) eq 'C') {
				my $TempMethStatus = substr ($Methylationsequence,14,1);
				$CGpos2=$cutcoord+12; $CGcand2 = uc(substr($sequence{$chr},$CGpos,4)); $seqmatch2 = reverse $CGcand2; $seqmatch2 =~ tr/ACGTacgt/TGCAtgca/; $CGcand2 = $seqmatch2; $strand2=-1; $NCHG++;

				if ($TempMethStatus eq 'X'){
				$CHGMaintained++;
				$NCHGMaintained="X";}
				elsif ($TempMethStatus eq 'x'){
				$CHGNotMaintained++;
				$NCHGMaintained="x";}
				else {$NCHGMaintained=".";}
			}
			
		}
		
		#Indirect, 5mC orignally on - strand
		if ($seqflag == 16 & substr($cutsite,96-$AdjustedLength,1) eq 'G'){
			$CGpos=$cutcoord+73-$AdjustedLength; $CGcand = uc(substr($sequence{$chr},$CGpos,4));$seqmatch = reverse $CGcand; $seqmatch =~ tr/ACGTacgt/TGCAtgca/; $CGcand = $seqmatch; $strand=-1; $Nhits++;
			}
			
		
		
		#note for last two 'G' CGpos is shifted by 3bp because of reverse complement !!    

		# if ($seqflag == 16){
		# for ($k=0;$k<105;$k++) {
		# my $TempNucleotide = substr($cutsite,$k,1);
							 
							 # if ($TempNucleotide eq "A") {$NucleotideA[$k]++;}
							 # if ($TempNucleotide eq "C") {$NucleotideC[$k]++;}
							 # if ($TempNucleotide eq "G") {$NucleotideG[$k]++;}
							 # if ($TempNucleotide eq "T") {$NucleotideT[$k]++;}
							 # }
							 # }


       if ($Nhits == 1 & $SpikeInFlag==0){print $abaout "$csbcindex\t$chrindex\t$CGpos\t$strand\t$umi\t$CGcand\t$Nhits\t$seqflag\t$NCGMaintained\t$NCHGMaintained\n";
$cleanreads[$csbcindex]++;}

		if ($NCG == 1 & $Nhits == 1 & $SpikeInFlag==0){print $CGabaout "$csbcindex\t$chrindex\t$CGpos2\t$strand2\t$umi\t$CGcand2\t$NCG\t$seqflag\t$NCGMaintained\n";}
		if ($NCHG == 1 & $Nhits == 1 & $SpikeInFlag==0){print $CHGabaout "$csbcindex\t$chrindex\t$CGpos2\t$strand2\t$umi\t$CGcand2\t$NCHG\t$seqflag\t$NCHGMaintained\n";}
	   if ($Nhits == 1 & $SpikeInFlag==1){print $Zymoabaout "$csbcindex\t$chr\t$CGpos2\t$strand2\t$umi\t$CGcand2\t$NCG\t$seqflag\t$NCGMaintained\n";}
	   if ($Nhits == 1 & $SpikeInFlag==2){print $Lamndaabaout "$csbcindex\t$chr\t$CGpos2\t$strand2\t$umi\t$CGcand2\t$NCG\t$seqflag\t$NCGMaintained\n";}

		}} 
print "$count1 $count2 $count3 $count4\n";
# write QC count to file
my $QCoutputfile = substr($file1,0,-3)."QC";
open(my $qcout, '>', $QCoutputfile) or die "Can't open file for writing: $!\n";
my $QCoutputfile2 = substr($file1,0,-3)."QC2";
open(my $qcout2, '>', $QCoutputfile2) or die "Can't open file for writing: $!\n";
for ($k=0;$k<96;$k++)
 {
   print $qcout "$k \t $rawreads[$k] \t $mappedreads[$k] \t $cleanreads[$k] \n";
   #print $qcout2 "$allreads[$k]\n";
 }

 print $qcout2 "A\tC\tG\tT\n";
			 for ($k=0;$k<105;$k++)
			 {
			   
			   print $qcout2 "$NucleotideA[$k]\t$NucleotideC[$k]\t$NucleotideG[$k]\t$NucleotideT[$k]\n";
			 }
 
 print "TotalUMINucleotides CUMINucleotides CGMaintained CGNotMaintained CHGMaintained CHGNotMaintained\n";

 print "$TotalUMINucleotides $CUMINucleotides $CGMaintained $CGNotMaintained $CHGMaintained $CHGNotMaintained\n";
# keep only unique cuts and sort
system("sort -u $rABAoutputfile > $fABAoutputfile");
system("sort -u $CGrABAoutputfile > $CGfABAoutputfile");
system("sort -u $CHGrABAoutputfile > $CHGfABAoutputfile");
system("sort -u $rABAoutputfileZymoSpikeIn > $fABAoutputfileZymoSpikeIn");
system("sort -u $rABAoutputfileLamndaPhageSpikeIn > $fABAoutputfileLamndaPhageSpikeIn");
```
