## Supplementary Software for "Combinatorial quantification of 5mC and 5hmC at individual CpG dyads and the transcriptome in single cells reveals modulators of DNA methylation maintenance fidelity": Pullout-scMaintainedCG_ForAbaSI.pl

```
#!/usr/bin/perl

use strict "subs";
use strict "refs";
#use strict "vars";
 

# read in faba
my $file1 = $ARGV[0] or die "Need to get CGfaba file on the command line\n";
open(my $data, '<', $file1) or die "Could not open '$file1' $!\n";

my $Maintainedoutputfile = substr($file1,0)."Maintained";

open(my $CGout, '>', $Maintainedoutputfile) or die "Can't open file for writing: $!\n";


for ($k=0;$k<96;$k++){ #Initialize counters per cell
	$CGMaintained[$k]=0;
	$CGNotMaintained[$k]=0;
	
	$CHGMaintained[$k]=0;
	$CHGNotMaintained[$k]=0;
}


while (my $line = <$data>) {
    chomp $line;
    my @fields = split "\t" , $line;
	my $cell = $fields[0];
    my $CGstatus = $fields[8];
	#my $CHGstatus = $fields[9];
	
		if ($CGstatus eq "z"){$CGNotMaintained[$cell]++;}
		elsif ($CGstatus eq "Z") {$CGMaintained[$cell]++;}
		#elsif ($CHGstatus eq "x"){$CHGNotMaintained[$cell]++;}
		#elsif ($CHGstatus eq "X"){$CHGMaintained[$cell]++;}
} 


for ($k=0;$k<96;$k++){
	$CGsum = $CGMaintained[$k]+$CGNotMaintained[$k];
	if($CGsum>0){$CGMaintenancePercent[$k]=$CGMaintained[$k]/$CGsum;}
	else{$CGMaintenancePercent[$k]="";}
	
	#$CHGsum = $CHGMaintained[$k]+$CHGNotMaintained[$k];
	#if($CHGsum>0){$CHGMaintenancePercent[$k]=$CHGMaintained[$k]/$CHGsum;}
	#else{$CHGMaintenancePercent[$k]="";}
	
	$cell = $k +1;
	print $CGout "$cell\t$CGMaintained[$k]\t$CGNotMaintained[$k]\t$CGMaintenancePercent[$k]\n";
}	
```
