## Supplementary Software for "Combinatorial quantification of 5mC and 5hmC at individual CpG dyads and the transcriptome in single cells reveals modulators of DNA methylation maintenance fidelity": Pullout-scMethylationFromBismarkMethylationExtractor_wReadNames.pl

```
#!/usr/bin/perl

use strict "subs";
use strict "refs";
#use strict "vars";
 

# read in faba
my $file1 = $ARGV[0] or die "Need to get MethylationExtractor on the command line\n";
open(my $data, '<', $file1) or die "Could not open '$file1' $!\n";

my $Maintainedoutputfile = substr($file1,0)."Format";

open(my $CGout, '>', $Maintainedoutputfile) or die "Can't open file for writing: $!\n";


my $BCIn = $ARGV[1];
open($BC, "$BCIn");


my @csbc;
#open(my $cscodes, '<', $BCIn) or die "Could not open '$file2' $!\n";
while (my $line = <$BC>) {
  chomp $line;
    my @fields = split "\t" , $line;
    push @csbc,@fields[0];
}
my $BARCODELength = length($csbc[0]);
my $UMILength = $ARGV[2] or die "Need UMI Length\n";

#File Name determines the strand + or -
$char = '/';
$NoPathIndex = rindex($file1, $char);
$NoPathName = substr($file1, $NoPathIndex+1);

$StrandVal = index($NoPathName,"CpG_OT_");
if ($StrandVal != 0){$StrandVal = index($NoPathName,"CpG_CTOT_");}
if ($StrandVal != 0){$StrandVal = index($NoPathName,"Non_CpG_OT_");}
if ($StrandVal != 0){$StrandVal = index($NoPathName,"Non_CpG_CTOT_");}

my @chrvec=qw(chr1 chr2 chr3 chr4 chr5 chr6 chr7 chr8 chr9 chr10 chr11 chr12 chr13 chr14 chr15 chr16 chr17 chr18 chr19 chr20 chr21 chr22 chrX chrY chrM ZymoDNAStandardSet PhageLambda);
while (my $line = <$data>) {
    chomp $line;
    my @fields = split "\t" , $line;
    my $cellbarcode = substr($fields[0],-1*$BARCODELength);
    my $chr = $fields[2];
    my $loci = $fields[3];
    my $UMI = substr($fields[0],-1*($BARCODELength+$UMILength),$UMILength);
	
    my ($csbcindex) = grep {$csbc[$_] ~~ $cellbarcode } 0 .. 95;  
    my ($chrindex) = grep {$chrvec[$_] ~~ $chr} 0 .. 26;

    my @ReadName = split "_", $fields[0];
    
	if ($cellbarcode ~~ @csbc & $chr ~~ @chrvec){
	    
	    if ($StrandVal == 0) {$strand = 1;}
	    else {$strand = -1;}
	    my $chrindex2 = $chrindex + 1;
	    my $csbcindex2 = $csbcindex + 1;
	    
	    print $CGout ("$csbcindex2\t$chrindex2\t$loci\t$strand\t$UMI\t$fields[4]\t$ReadName[0]\n");
	}
} 
```
