## Supplementary Software for "Combinatorial quantification of 5mC and 5hmC at individual CpG dyads and the transcriptome in single cells reveals modulators of DNA methylation maintenance fidelity": ReadMe.txt

Refer to manuscript for computation pipeline description. A step by step guide and accompanying shell scripts and additional information to run all code sequentially can be found at https://github.com/alexchialastri/scDyad-T-seq

## useage of each script below ##

5mCpGDiadSingleCellBarcodes.txt - is the list of scDyad-seq barcodes described in the manuscript

AbaSI_5hmC_CpGDyad_SingleCell_Barcodes_First3.txt - is the list of barcodes described in the mansucript for AbaSI based Dyad-seq

AbaSIFabaToCGtxt.pl - Takes .faba file and creates a CpG hydroxymaintained (fully hydroxymethylated) and non-maintained files (hemi-hydroxymethylated) text files. These resulting files are used in downstream analysis and are submitted on GEO.
Argument 0 = .faba file generated by the process_sc5hmC_CpGDiad.pl script.

FabaToCGtxt.pl - Takes .faba file and creates a CpG maintained (fully methylated) and non-maintained files (hemi-methylated) text files. The exact loci of the CpG site is modified slightly from the faba file to correct for which strand it was detected on. These resulting files are used in downstream analysis and are submitted on GEO.
Argument 0 = .faba file generated by the process_scCpGDiadwithQC.pl script.

MakeFastqShorter.pl - Used to hard clip Fastq files
Argument 0 = Input Fastq file name
Argument 1 = Desired Output Fastq file name
Argument 2 = Desired Fastq file trimmed length

process_sc5hmC_CpGDiad.pl - Searches the genome location of mapped reads to identify fully or hemi-hydroxymethylated CpG sites. Also detects all 5hmC calls inferred by the cutting action of AbaSI.
Argument 0 = Genome used for mapping (non bisulfite converted)
Argument 1 = Mapped deduplicated sam file from Bismark
Argument 2 = Cell barcodes used for H-M-Dyad-seq, H-H-Dyad-seq.
Argument 3 = UMI length used (typically = 4)

process_scCpGDiadwithQC.pl - Searches the genome location of mapped reads to identify fully or hemi-methylated CpG or CpHpG sites. Also detects all 5mC calls inferred by the cutting action of MspJI.
Argument 0 = Genome used for mapping (non bisulfite converted)
Argument 1 = Mapped deduplicated sam file from Bismark
Argument 2 = Cell barcodes used for M-M-Dyad-seq, M-H-Dyad-seq or scDyad-seq.
Argument 3 = UMI length used (typically = 4)

Pullout-scMaintainedCG_ForAbaSI.pl - Summarizes the detection per cell. Resulting .fabaMaintained text file gives the cell number, the number of fully hydroxymethylated detections, the number of hemi-hydroxymethylated detections, and the percent fully hydroxymethylated for CpG sites.
Argument 0 = .faba file generated by the process_sc5hmC_CpGDiad.pl script.

Pullout-scMaintainedCGandCHG.pl - Summarizes the detection per cell. Resulting .fabaMaintained text file gives the cell number, the number of fully methylated detections, the number of hemimethylated detections, and the percent fully methylated for CpG sites followed by CpHpG sites in the next 3 columns.
Argument 0 = .faba file generated by the process_scCpGDiadwithQC.pl script.

Pullout-scMethylationFromBismarkMethylationExtractor_wReadNames.pl - Reformats the Bismark Methylation Extractor format to demultiplex by cell barcode and UMI.
Argument 0 = Bismark Methylation Extractor text file. There are 8 options. [CpG_OB_*txt, CpG_CTOB_*txt, CpG_OT_*txt, CpG_CTOT_*txt, Non_CpG_OB_*txt, Non_CpG_CTOB_*txt, Non_CpG_OT_*txt, Non_CpG_CTOT_*txt]. Run this script on each of the Bismark Methylation Extractor output files.
Argument 1 = Barcode list
Argument 2 = UMI length (typically = 4)

scCpGDiad_Trim_and_ExtractBarcodes_se-HammingCorrection_TransferToR2.pl - Used to identify reads with the proper PCR sequence "GGTGTAGTGGGTTTGG" and proper barcodes. Also concatenates the UMI and cell barcode to the read name for read 1 and 2
Argument 0 = Barcode list
Argument 1 = Read 1 Input Fastq file
Argument 2 = UMI length (typically = 4)
Argument 3 = Read 2 Input Fastq file


 
