## Supplementary Software for "Combinatorial quantification of 5mC and 5hmC at individual CpG dyads and the transcriptome in single cells reveals modulators of DNA methylation maintenance fidelity": scCpGDiad_Trim_and_ExtractBarcodes_se-HammingCorrection_TransferToR2.pl

```
# Extract CpG Diad reads based on 96 barcodes from fastq files (containing UMIs too) 
# Also has to match the GGTGTAGTGGGTTTGG PCR adpt
# Also puts the Barcode and UMI infront of the Read Name followedby a -                                                                                                         
use warnings;
my $start_run = time();

my $fastafileR1In = $ARGV[1];

open($fastafileR1, "$fastafileR1In");

my $fastafileR2In = $ARGV[3];
open($fastafileR2, "$fastafileR2In");


my $outputR1In = substr($fastafileR1In,0,-6)."-BS_of_CpGDiad.fastq";
my $outputR2In = substr($fastafileR2In,0,-6)."-BS_of_CpGDiad.fastq";


open($outputR1, '>', "$outputR1In");
open($outputR2, '>', "$outputR2In");

my $BCIn = $ARGV[0];
open($BC, "$BCIn");

$PCRCode="GGTGTAGTGGGTTTGG";
$PCRCodeLength = length($PCRCode);

$UMILength=$ARGV[2]; #Length of the UMI barcode


$i = 0;
while ($BCline = <$BC>) #read in barcodes
{
    chomp($BCline);
    $BarCode[$i] = $BCline;
    $i++;                                                                                                                                                    
    
}
$BarcodeLength= length($BarCode[0]); #Length of the Cell barcode
                             

while ($header = <$fastafileR1>)
{
    chomp ($header);    
    $sequence = <$fastafileR1>;
    $plus     = <$fastafileR1>;
    $qual     = <$fastafileR1>;


    $headerR2 = <$fastafileR2>;
    chomp ($headerR2);    
    $sequenceR2 = <$fastafileR2>;
    $plusR2     = <$fastafileR2>;
    $qualR2     = <$fastafileR2>;


 $MatchFlag=0;
$SequencedCellBarcode = substr($sequence,$PCRCodeLength+$UMILength,$BarcodeLength); #20 because PCR barcode is 16 long and the UMI is 4 [DWDD]

    for ($i=0;$i<96;$i++)
    { #Cycle through barcodes
	
	
	$start1a = 0;
	for ($k=0;$k<$BarcodeLength;$k++){ #go base by base in the barcode to see how many match +1 for each matching base
		$TempBarcodeSequencedBase = substr($SequencedCellBarcode,$k,1);
		$TempBarcodeRightBase = substr($BarCode[$i],$k,1);
		    if ($TempBarcodeSequencedBase eq $TempBarcodeRightBase){$start1a++;}
	    }

	if ($start1a >= $BarcodeLength-1)
	{
	    #Correct Barcode Sequence for those off by 1 base
	    $UMIBC = substr($sequence,$PCRCodeLength,$UMILength); #UMI is 4 Long
	    $header2 = $header . '-:' . $UMIBC.$BarCode[$i];
	    $sequence2=substr($sequence,$PCRCodeLength+$UMILength+$BarcodeLength);
	    $qual2=substr($qual,$PCRCodeLength+$UMILength+$BarcodeLength);
	    
	    $headerR22 = $headerR2 . '-:' . $UMIBC.$BarCode[$i];

	    #I want to check if the PCR Barcode matches as well just to be sure but
	    #Allow Some Mismatch in PCR Barcode
	    $start1b = 0;
	    $SequencedPCRCode = substr($sequence,0,$PCRCodeLength);
	    for ($j=0;$j<16;$j++){#same concept as done with cell barcode
		$TempPCRSequencedBase = substr($SequencedPCRCode,$j,1);
		$TempPCRRightBase = substr($PCRCode,$j,1);
		    if ($TempPCRSequencedBase eq $TempPCRRightBase){$start1b++;}
	    }
	    
		
		if ($start1b >= 15)
		{#If within 1 hamming of a cell barcode and 1 of the PCR barcode, reprint it here with the Header Line - UMI Cell BC
		
		$MatchFlag=1;
		#chomp($header2);
		#chomp($sequence2);
		#chomp($plus);
		#chomp($qual2);
		#print $outputR1 ("$header2\n$sequence2\n$plus\n$qual2\n");
		print $outputR1 ("$header2\n$sequence2$plus$qual2");
		print $outputR2 ("$headerR22\n$sequenceR2$plusR2$qualR2");
		last;
		}
		

	}
    }
#if ($MatchFlag==0) {print $output2R1 ("$header\n$sequence$plus$qual");}
}

my $end_run = time();
my $run_time = $end_run - $start_run;
print "Job took $run_time seconds\n";
```
